# Long-Read epigenetic clocks identify improved brain aging predictions

**DOI:** 10.1101/2025.09.30.679553

**Authors:** Spencer M. Grant, Sarah J. Eger, Mary B. Makarious, Melissa Meredith, Abraham Moller, Melissa Grant-Peters, Amy Hicks, Ajeet Mandal, Pavan Auluck, Andrés Villegas-Lanau, Bárbara Mejia-Cupajita, Juliana Acosta-Uribe, David Aguillón, Hampton Leonard, Nicole Kuznetsov, Cory Weller, Adam Catching, Xylena Reed, Miten Jain, Luigi Ferrucci, Kenneth S. Kosik, Mark R. Cookson, Mina Ryten, Mike A. Nalls, Kimberley J. Billingsley

## Abstract

Epigenetic clocks are widely used to estimate biological aging, yet most are built from array-based data from peripheral tissues of predominantly European-ancestry individuals, limiting their generalizability. Here, we present aging clocks on DNA methylation from Oxford Nanopore long-read sequencing (LRS), leveraging over 28 million CpG sites from prefrontal cortex samples across individuals of African and European ancestry. These models were developed using GenoML, an automated machine learning platform for multi-omics data that leverages a diverse catalog of existing model architectures. Our long-read-informed clocks were developed using promoter-based and whole-genome window-based features, yielding models for each individual cohort as well as a combined-cohort clock. Each of these models demonstrated favorable performance compared to existing methylation clocks and was externally validated in a cohort of Colombian individuals. We further performed enrichment analyses and nominated both shared and cohort-specific pathways, cell types, and transcription factor binding motifs which may be implicated in aging and were not fully explained by cell type proportions or postmortem interval. Altogether, our findings highlight the power of long-read methylation data for constructing accurate, ancestry-aware aging clocks and emphasize the importance of inclusive training datasets.

## Introduction

DNA methylation serves as a uniquely valuable modality for assessing biological aging due to its predictable age-associated patterns and its reflection of both genetic and environmental factors ^1^. Consequently, epigenetic clocks, which are machine learning (ML) models that translate methylation data to determinations of biological age, have become increasingly relevant in tracking biological mechanisms associated with aging ^1,2^.

The first methylation clock, published by Steve Horvath in 2013, successfully associated methylation with aging across diverse tissues and cell types ^1^. Since then, the concept of utilizing DNA methylation as a predictor for health indicators has expanded both in scope and utility. This includes second-generation (to predict healthspan) ^3,4^, third-generation (to predict pace of aging) ^5^, and fourth-generation (to identify causal underpinnings of aging) ^6^ clocks ^7^. As most of these clocks use individual CpG sites as the primary predictive methylation features, principal component analysis (PCA)-based models have gained traction to alleviate potential stochasticity ^8^ associated with isolated CpGs while reducing the dimensionality of the datasets ^9^. PCA models provide a more stable alternative to CpG-based models, albeit with reduced interpretability. While the landmark Horvath clock was robust across diverse tissues and cell types, it was developed on human samples of predominantly European ancestry. Since then, there have been more recent efforts to further expand this across mammalian species, which included samples from non-European individuals, using highly-conserved genomic regions ^2^.

Most methylation clocks have been developed using variations of linear regression models, such as Elastic Net ^2,10^, which form an optimized linear combination of features. These ML models are well-suited for epigenetic clocks due to their ability to combine DNA methylation reads across the genome, providing insights about an individual’s biological age based on molecular features. One potential shortcoming of linear ML architectures in modeling aging, however, is their inability to capture the well-established nonlinearity in aging mechanisms ^11,12^. Deep learning ^13^ is theoretically suitable for handling such nonlinear relationships and complex feature interactions; however, it requires substantially larger datasets to generalize, with minimal evidence of superiority over linear models in practice ^14^.

Existing methylation clocks, however, have several limitations. Most clocks have been generated using array-based datasets, which capture less than 5% of CpG sites genome-wide, thus restricting the depth of the epigenomic feature sets available to be used by ML models. To address this, recent advances in long-read sequencing now enable high-resolution methylation profiling across the genome. Unlike array- or bisulfite-based methods, Oxford Nanopore Technologies (ONT) long-read sequencing provides genome-wide methylation data at single-molecule resolution. At the *APOE* locus, which bears genetic risk across multiple neurodegenerative diseases ^15,16^ with ancestry-specific genomic characteristics ^17^, ONT captures 33 times more CpGs than the Illumina 450K array, offering greater insight into densely methylated and functionally relevant regions ^18^. Additionally, many existing methylation clocks exhibit varied results when applied to samples from non-European individuals ^19^ due to their underrepresentation in datasets used to train their models. This reflects the principle that ML model generalizability depends on training data characteristics. Relatedly, tissue-specific clocks have been shown to be advantageous to blood-based and even multi-tissue clocks ^20–23^, including in brain tissue ^24,25^; however, the availability of brain-specific models and datasets remains relatively sparse compared to those derived from blood.

Here, we present methylation clocks that have been developed from long-read DNA methylation sequencing datasets in prefrontal cortex (PFC) samples from both European and African/admixed-ancestry individuals. Because age is the predominant risk factor for neurodegenerative disorders, brain samples are valuable targets for both epigenetic age modeling and mechanistic insight, with distinct methylation characteristics from peripheral tissues ^26^. Rather than relying on individual CpG sites, which may be prone to technical variability, or PCA features, which lack interpretability, we developed region-based features by aggregating signal over promoters and genome-wide windows. Our long-read-informed clocks (LR-clocks) were produced using GenoML, an automated ML tool that competitively selects the best-performing ML architecture from a pool of both linear and nonlinear algorithms ^27^. These measures yielded models which performed favorably compared to benchmark clocks while also preserving biological interpretability. In addition to cohort-specific clocks, we generated combined-cohort clocks that performed comparably in each cohort. To discern brain aging mechanisms, we identified the most important genes and promoters in each of our LR-clocks to serve as the basis for enrichment analyses, nominating several pathways, cell types, and transcription factor (TF) binding motifs which may be differentially important in brain aging across cohorts.

## Results

### Data overview

The data used in this study include long-read methylation reads from PFC tissue from the North American Brain Expression Consortium (NABEC, USA) and the Human Brain Collection Core (HBCC, USA). The NABEC cohort includes harmonized methylation and clinical data from 189 individuals of European ancestry and the HBCC cohort includes harmonized data from 132 individuals of African or admixed ancestry. All participants were neurologically healthy, making these cohorts suitable for ancestry-specific methylation clocks. As shown in **Figure 1**, the workflow includes long-read whole-genome sequencing, aggregation of DNA methylation signal across genomic regions, and predictive modeling using GenoML ^27^, which utilizes a competitive approach for algorithm selection that is designed to account for the unique structures and complexities long-read methylation datasets. More details about GenoML and the cohorts included here can be found in the Methods.

**Figure 1:**
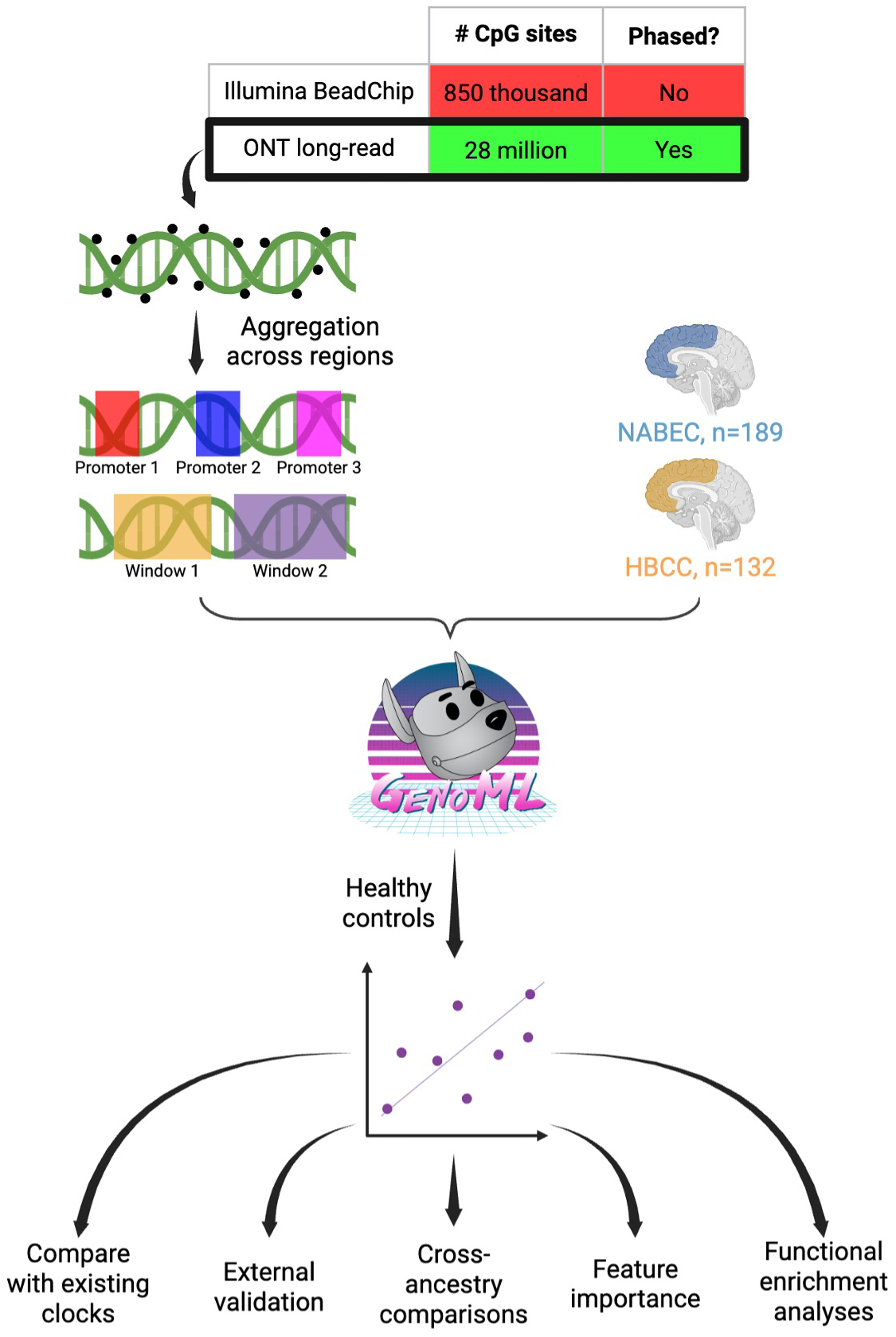
Long-read methylation and machine learning workflow. Schematic overview of the data generation and analysis pipeline, from long-read WGS through region-based DNA methylation aggregation, predictive modeling using *GenoML*, benchmarking against existing methylation clocks, external validation, and downstream functional enrichment analyses.

### LR-clocks demonstrate favorable performance compared to existing clocks

To establish baseline models, we developed methylation clocks based on the CpG sites used in several established methylation clocks that span a diverse spectrum of intended use cases. These include the first-generation Horvath pan-tissue clock (334 CpGs from existing arrays) ^1^; the second-generation ancestrally diverse PhenoAge clock (513 CpGs) ^3^; the third-generation DunedinPACE clock (173 CpGs) ^5^; the three pan-mammalian, pan-tissue clocks from Lu *et al.* (335, 816, and 760 CpGs, respectively) ^2^; and the brain-specific clock from Shireby *et al.* (347 CpGs) ^24^. We first generated one model for each benchmark clock for each sample set (each cohort individually, plus one combined-cohort set). This yielded mixed results, with consistently more accurate predictions on participants from the NABEC cohort compared to HBCC across all benchmark models tested **(Supplementary Figures 1-3, Supplementary Tables 1-3)**. The NABEC-specific clocks likewise moderately outperformed each of the combined-cohort clocks, which themselves outperformed each of the HBCC-specific clocks.

We then conducted a second benchmark analysis, similarly focused on the same loci from each benchmark clock, but using the aggregated methylation signal throughout the nearest promoter (see Methods for details about the EPDnew promoters used here) to each locus in lieu of individual CpGs. This aggregation step aimed to reduce stochastic variability while preserving each region’s functional significance. These promoter-based models performed substantially better, exhibiting improved accuracy for nearly all benchmark models across all sample sets **(Figure 2**, **Table 1)**. The improved accuracy in the promoter-based models from the CpG-based models was especially pronounced in HBCC, which performed much more comparably to the other sample sets.

**Figure 2.**
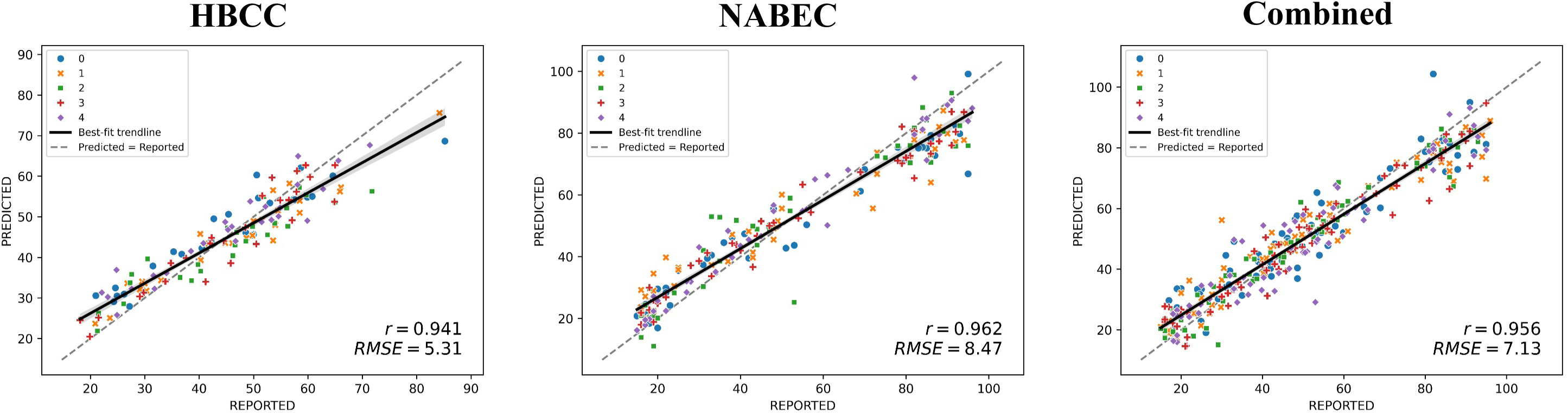
Long-read promoter-based clock results. Scatterplots depicting predicted age vs. reported age for each sample across all five outer cross-validation folds. The solid line depicts the best-fit trendline to the data, while the dashed line represents the points where the predicted age is equal to the reported age. The three scatterplots correspond to the promoter-based long-read clocks for the three sample sets – HBCC-only, NABEC-only, and HBCC and NABEC combined, for both training and testing.

**Table 1:** Accuracy metrics for promoter-based long-read clocks.

|  | Explained Variance | Root Mean Squared Error | Median Absolute Error | Pearson's R |
| --- | --- | --- | --- | --- |
| HBCC | 0.869 | 5.274 | 3.519 | 0.949 |
| NABEC | 0.906 | 8.368 | 5.747 | 0.965 |
| Combined-cohort | 0.906 | 7.073 | 4.194 | 0.956 |

Finally, using a similar methodology to the promoter-based regions, we aggregated signal across whole-genome windows, subsetting to include the set of windows from the long-read data that overlap with the CpG set from each respective benchmark clock. Genomic coordinates for each whole-genome window can be found in **Supplementary Table 4**. In addition to reducing noise-related overfitting, this also aimed to more fully leverage genome-wide resolution provided by long-read sequencing. With a few exceptions, the window-based models consistently yielded improvements over the promoter-based models across all sample sets **(Figure 3**, **Table 2)**. While the Horvath and DunedinPACE benchmarks routinely produced weaker results in the HBCC cohort, results were generally comparable across all sample sets for all other benchmark clocks using whole-genome window features.

**Figure 3.**
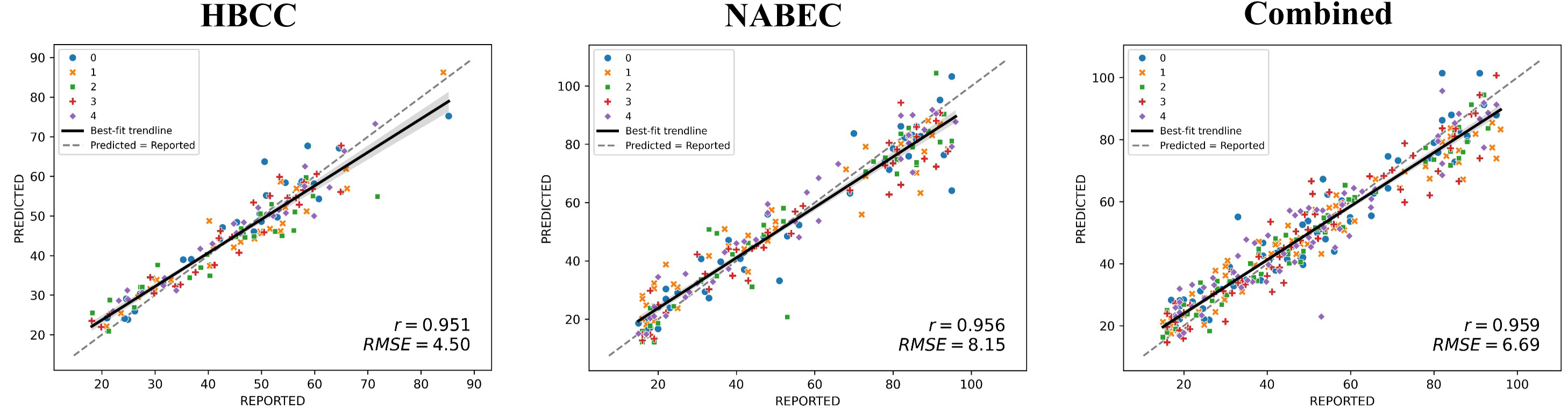
Long-read window-based clock results. Scatterplots depicting predicted age vs. reported age for each sample across all five outer cross-validation folds. The solid line depicts the best-fit trendline to the data, while the dashed line represents the points where the predicted age is equal to the reported age. The three scatterplots correspond to the whole-genome window-based long-read clocks for the three sample sets – HBCC-only, NABEC-only, and HBCC and NABEC combined, for both training and testing.

**Table 2:** Accuracy metrics for window-based long-read clocks.

|  | Explained Variance | Root Mean Squared Error | Median Absolute Error | Pearson's R |
| --- | --- | --- | --- | --- |
| HBCC | 0.906 | 4.426 | 3.066 | 0.958 |
| NABEC | 0.915 | 8.090 | 4.484 | 0.96 |
| Combined-cohort | 0.918 | 6.632 | 3.878 | 0.961 |

After establishing the predictive accuracy of long-read methylation data using previously reported CpG sites, we sought to create methylation clocks based on unbiased identification of genomic regions within each sample set. We first performed CpG-level quality control (QC), which is described in detail in the Methods. Starting with all 29,598 promoters included in the Eukaryotic Promoter Database ^28^ and the 559,368 whole-genome windows as described in the Methods, filtering steps were performed separately within each sample set to remove features that are highly correlated with each other and to keep only those which are significantly predictive of age. This yielded 7,337; 5,693; and 11,099 promoters (of which 368; 5,631; and 1,338 contributed to the performance of the final models) and 109,542; 146,108; and 159,972 windows (of which 1,834; 2,990; and 2,039 contributed to the performance of the final models) which were used as the final feature set for the HBCC, NABEC, and combined-cohort sample sets, respectively. These LR-clocks predicted chronological age with high accuracy in each sample set (promoter-based: HBCC R^2^ = 0.862, NABEC R^2^ = 0.905, Combined R^2^ = 0.905; window-based: HBCC R^2^ = 0.901, NABEC R^2^ = 0.912, Combined R^2^ = 0.917), producing more accurate predictions than the benchmarking models **(Supplementary Figures 1-3, Supplementary Tables 1-3)**. Intercepts and beta coefficients from each of the LR-clocks can be found in **Supplementary Tables 5-10.** A summary of the overlap between each of the promoter-based LR-clocks and each of the benchmark clocks can be found in **Table 3**.

**Table 3:** Summary of overlap between promoters used in the final long-read models and nearest promoters to clock sites taken from existing clocks used for benchmarking. “Within” represents CpG sites that are within overlapping promoters, while “adjacent” represents CpG sites that are not within overlapping promoters, for which the nearest promoter was used instead.

|  | **HBCC (Total = 368 promoters)** | **NABEC (Total = 5693 promoters)** | **Combined (Total = 1338 promoters)** |
| --- | --- | --- | --- |
| **DunedinPACE (Total = 173 CpGs)** | 3 (0 within, 3 adjacent) | 37 (12 within, 25 adjacent) | 10 (5 within, 5 adjacent) |
| **Horvath (Total = 334 CpGs)** | 4 (4 within, 0 adjacent) | 61 (53 within, 8 adjacent) | 23 (21 within, 2 adjacent) |
| **Lu_Clock1 (Total = 335 CpGs)** | 20 (8 within, 12 adjacent) | 90 (29 within, 61 adjacent) | 35 (14 within, 21 adjacent) |
| **Lu_Clock2 (Total = 816 CpGs)** | 38 (11 within, 27 adjacent) | 227 (56 within, 171 adjacent) | 83 (26 within, 57 adjacent) |
| **Lu_Clock3 (Total = 760 CpGs)** | 33 (9 within, 24 adjacent) | 199 (54 within, 145 adjacent) | 74 (21 within, 53 adjacent) |
| **PhenoAge (Total = 513 CpGs)** | 5 (4 within, 1 adjacent) | 110 (86 within, 24 adjacent) | 31 (25 within, 6 adjacent) |
| **Shireby (Total = 347 CpGs)** | 14 (8 within, 6 adjacent) | 110 (59 within, 51 adjacent) | 48 (28 within, 20 adjacent) |

Across all tested models, the Shireby brain-specific clocks uniformly outperformed each of the other benchmark clocks. The promoter-based LR-clocks yielded favorable results compared to the Shireby clocks across all sample sets. For the window-based clocks, the results were much more comparable, with the Shireby clock showing slightly higher accuracy in HBCC while the LR-clocks resulted in higher accuracy in NABEC and the combined-cohort set. In general, across all benchmark and LR-clocks, models developed on whole-genome windows performed favorably to those developed on promoter regions, which in turn outperformed corresponding CpG-based models. These findings indicate that (a) brain-specific models outperform generalized multi-tissue models for brain aging predictions; (b) regional signal aggregation provides valuable context into aging mechanisms; and (c) the inclusion of more features genome-wide improves ML model performance.

Due to the expanding evidence indicating the effectiveness of methylation principal components (PCs) as features ^9^, we developed PC-based clocks on the CpG-level long-read data for each sample set after removing low-variance CpGs. PCs were then used as input features for LR-clocks, which uniformly resulted in weaker correlations than each of the other LR-clock feature sets **(Supplementary Figures 1-3, Supplementary Tables 1-3)**, supporting the robustness of signal aggregation across promoters and whole-genome windows for dimensionality reduction. As an additional benefit of direct signal aggregation, these methods preserve biological interpretability while the physiological meaning of principal components is much more difficult to ascertain.

Of the 72 total clocks that resulted from this workflow, GenoML identified ElasticNet as the optimal architecture in all but eight. These included five instances of Gradient Boosting Regressor (combined-cohort DunedinPACE promoters, combined-cohort DunedinPACE windows, combined-cohort PCA LR-clock, combined-cohort Lu Clock 1 windows, and NABEC PCA LR-clock), two of AdaBoost Regressor (NABEC and HBCC PCA LR-clocks), and one of K-Neighbors Regressor (HBCC DunedinPACE CpGs).

### Covariates do not fully explain observed age acceleration

To assess whether age acceleration estimates derived from our LR-clocks were driven by known confounds, we examined associations between age acceleration and both post-mortem interval (PMI) and estimated cell type composition in each sample set. Age acceleration was defined as the residual from regressing predicted age on chronological age. PMI was not significantly associated with age acceleration in any sample set individually or jointly with cell type composition (all FDR-corrected p > 0.05), with negligible effect sizes (β = -0.028 to 0.037 years per hour). Cell type composition, which has been shown to be modestly correlated with chronological age ^29^, collectively explained 1.5-6.9% of variance in age acceleration across sample sets, with no individual cell type surviving FDR correction. However, inhibitory neuron fraction showed a consistent negative association with age acceleration across all three sample sets and nominal significance in the HBCC joint cell type + PMI model (**Supplementary Tables 11-13, Supplementary Figure 5**), consistent with known inhibitory neuron vulnerability in aging ^30^ and several neurodegenerative diseases ^31,32^. This suggests these variables largely track age-related changes already captured by the clock’s chronological age prediction, rather than driving residual aging-rate variation. Together, these results support that individual aging-rate variation captured by our clocks predominantly reflects genuine biological signals rather than sampling artifacts.

### LR-clock feature importance nominates shared and cohort-specific aging-relevant regions

Feature importance plots were generated using the Shapley Additive exPlanation (SHAP) package, which ranks features by their overall contributions to model predictions. The top 20 features for each LR-clock are visualized in **Supplementary Figure 6**, with a comprehensive list of SHAP values available in **Supplementary Tables 14-19**. Interestingly, the features with the maximum |SHAP| values for both the promoter-based and window-based HBCC models are larger than those of their NABEC and combined-cohort counterparts, while the sum of all |SHAP| values is smaller. This reflects the use of fewer features in the HBCC clocks with nonzero SHAP values (368 and 1,834, respectively) than in the NABEC (5,693 and 2,990, respectively) and combined-cohort (1,338 and 2,039, respectively) clocks, particularly in the promoter-based models.

Several specific EPDnew promoter features appeared in the top 20 in at least two of the promoter-based clocks. These included MLNR_1 (NABEC and HBCC), NETO2_1 (combined-cohort and HBCC), ANK1_3 (combined-cohort and HBCC), HHEX_1 (combined-cohort and HBCC), BNC1_1 (combined-cohort and NABEC), and CCNI2_1 (combined-cohort and NABEC). Each of these, with the exception of CCNI2_1, was included in all three models and had higher methylation associated with older predicted age in each model in which it was retained. Additionally, two different EPDnew promoter features assigned to *HOXD8* appeared in the top 20 for the combined-cohort (HOXD8_2) and the HBCC (HOXD8_1) models, with both also contributing to the NABEC model, all with positive SHAP values. Notably, and consistent with results reported elsewhere ^33^, several EPDnew promoter features assigned to genes within protocadherin (PCDH) clusters had nonzero SHAP values in the NABEC and combined-cohort models, with higher methylation at almost all of these features associated with an older predicted ages. Interestingly, only one PCDH promoter contributed to the HBCC promoter-based model, with higher methylation associated with a small shift toward a younger predicted age.

Several of these recurrent promoters are of particular biological interest. *ANK1* encodes ankyrin-1, a scaffolding protein that organizes membrane domains in neurons and has been previously implicated in Alzheimer’s disease ^34–36^, suggesting it may be reflective of preclinical neurodegeneration ^34^. *NETO2* encodes an auxiliary subunit of kainate receptors that regulates both excitatory synaptic transmission and chloride homeostasis through KCC2 modulation, the decline of which is a documented feature of neuronal aging ^37^. *HHEX* is a transcriptional repressor involved in embryonic forebrain development ^38^ and suppression of axon regeneration in mature neurons ^39^. This may reflect the progressive dysregulation of developmental transcriptional programs with age.

For the window-based models, three whole-genome windows appeared in the top 20 in multiple clocks: windows 248075 (HBCC and NABEC, overlapping the promoter region represented by ANK1_3), 196025 (HBCC and NABEC, overlapping the promoter region represented by SIM1_1), and 16825 (combined-cohort and HBCC). ANK1_3, as discussed above, was a strong contributor across each sample set, while SIM1_1 was among the top 20 features in the HBCC promoter-based model and contributed to both the NABEC and combined-cohort promoter-based models. Each of these whole-genome windows has a nonzero value in all window-based LR-clocks. With the exception of window 16825, for which higher methylation shifted predictions toward a younger age in the NABEC model, higher methylation at each window was associated with an older predicted age across the LR-clocks. Taken together with the aforementioned promoters, this identifies recurrent age-predictive loci captured using both the promoter-based and window-based aggregation methods.

### Functional enrichment analyses reveal shared and cohort-dependent biological mechanisms of brain aging

From each of the LR-clocks, we identified genes corresponding to promoters or windows with nonzero SHAP values in their respective models, as well as promoters overlapping with each whole-genome window. These were used as the basis for functional enrichment analyses, which were performed separately for promoters/genes associated with promoters/windows that are hypermethylated vs. hypomethylated with age. To avoid redundancy, genes associated with multiple promoters or windows were included only once.

#### HOMER implicates transcriptional regulators of aging-associated methylation changes

Hypergeometric Optimization of Motif EnRichment (HOMER) was used to identify transcription factor (TF)-binding motifs enriched in promoter sets associated with negatively and positively age-associated methylation features in each LR-clock. The top five known and *de novo* TF motifs for each set of promoters are shown in **Supplementary Figures 7-10**. Consistent with prior literature ^40^, we identified REST as a significantly-enriched motif in hypermethylated promoters, while also observing Arnt and CTCF (Zf) as being enriched in the hypomethylated promoters from the promoter-based combined-cohort and NABEC models, respectively. Interestingly, the REST motif was identified as the most significantly-enriched motif in each of the NABEC and the combined-cohort promoter-based and window-based models, while not being identified by either HBCC model altogether. HOMER also identified numerous zinc finger transcription factors across all sample sets in both the hypo- and hyper-methylated promoter sets.

There were, additionally, several motifs that recurred across more than one model and were annotated to factors with potential relevance to brain aging. BACH1, enriched in hypomethylated promoters in the NABEC and combined-cohort models, recognises Maf response elements also bound by NRF2-containing complexes ^41^. This finding nominates oxidative-stress-responsive regulatory sequences, although it does not establish which factor occupies them or the direction of transcriptional regulation. ZBTB38, enriched in hypermethylated regions in both HBCC and NABEC, is a methyl-CpG binding protein ^42^, indicating age-related methylation at these sites may recruit ZBTB38 to propagate downstream transcriptional silencing. CDX4, a developmental patterning factor that cooperates with HOX genes in neural progenitor specification ^43^, was enriched in hypermethylated promoters across HBCC and NABEC models, consistent with an overrepresentation of developmental regulatory sequences among these features. We additionally observe concordance across multiple models supporting the significance of Nr5a2 (hypomethylated in HBCC promoter-based and NABEC window-based models), NF1 (hypomethylated in NABEC promoter-based and combined-cohort window-based models), and Zfp335 (hypomethylated in HBCC and combined-cohort window-based models).

#### Expression-weighted cell type enrichment identifies distinct cell types epigenetically associated with brain aging

Using expression-weighted cell type enrichment (EWCE), we tested whether genes mapped to age-associated methylation features were preferentially expressed in particular brain cell types. Across all three sample sets, genes preferentially expressed in astrocytes were enriched among genes mapped to hypomethylated window features. Similarly, genes preferentially expressed in oligodendrocyte progenitor cells (OPCs) were enriched among genes mapped to hypermethylated window features in all three sample sets. While these cell types showed strong concordance, other enrichment patterns differed between cohorts. The NABEC promoter-based model revealed broad and significant enrichment across nearly all brain cell types for both hyper- and hypo-methylated promoters, while the HBCC promoter-based model yielded no significant enrichment in any cell type. The combined-cohort promoter-based model identified superficial excitatory neurons (L2/3) among hypermethylated genes and astrocytes among hypomethylated genes. Window-based models revealed more complex patterns. Among hypermethylated genes, NABEC and the combined cohort showed broad enrichment across excitatory and inhibitory neuron subtypes and glial populations, while HBCC showed no significant enrichment. Among hypomethylated window genes, enrichment was more concordant between HBCC and the combined cohort, with both showing enrichment for genes preferentially expressed in excitatory neuronal cell types. In contrast, NABEC showed enrichment for genes preferentially expressed in inhibitory neuronal, endothelial/mural, OPC, and microglial cell types.

#### Gene ontology analysis reveals pathway-level mechanisms of age-associated genes across cohorts

Gene ontology (GO) revealed several pathway-level insights about the LR-clocks (**Supplementary Figure 11, Supplementary Tables 20-23**). Many of the significantly enriched GO terms across each of the four sets of genes (hypo-vs. hyper-methylated, window-vs. promoter-based models) did not overlap between the HBCC and NABEC cohorts, indicating that the sets of nominated functional annotations differed between the cohorts. Similarly, we observe substantial differences in the GO pathways identified between the promoter-based and window-based models. This likely reflects a fundamental difference in scope – promoter features capture methylation signal restricted to annotated promoter regions, whereas window features tile the genome in non-overlapping 1kb windows, capturing methylation across a broader set of genomic contexts including gene bodies, intergenic regions, and non-promoter regulatory elements (in addition to promoters). The two approaches may therefore nominate complementary sets of functional annotations, with promoter-based models restricted to annotated transcription start-site regions and window-based models sampling a broader genomic context ^44,45^. Furthermore, several pathways were found to be significant in the combined-cohort models but not in either of the individual-cohort models. This may reflect generalized aging mechanisms that are captured by models trained on diverse cohorts that would otherwise be diluted by cohort-specific patterns, highlighting the importance of utilizing diverse samples when developing methylation clocks. Additionally, there were altogether fewer pathways identified using hypomethylated genes compared to hypermethylated genes, suggesting hypermethylated genes show stronger pathway-level coordination. This disparity is particularly true with the promoter-based models, with window-based models nominating a comparable number of pathways from the hypo-vs. hyper-methylated gene sets, in line with previous literature demonstrating general patterns of global hypomethylation and promoter hypermethylation with age ^46,47^. This is also consistent with the findings from HOMER, in which p-values for the most significant TF-binding motifs were consistently weaker for hypomethylated regions than for hypermethylated regions.

More specifically, GO results identified different functional themes among genes mapped to age-associated methylation features across cohorts, highlighted by a notable cohort-specific divergence in hypermethylated regions. Specifically, hypermethylated age-associated genes identified in the HBCC window-based model were strongly enriched for neuronal processes. Meanwhile, NABEC exhibited stronger enrichment for broader developmental and morphogenetic pathways, such as pattern specification and skeletal system development, primarily in the promoter-based model. Hypermethylated genes across both cohorts for both promoter- and window-based clocks consistently shared molecular functions related to sequence-specific DNA binding, transcription factor activity, and RNA polymerase binding. Conversely, hypomethylated genes in the combined-cohort models demonstrated enrichment for anterior-posterior pattern specification and embryonic morphogenesis.

### LR-clocks exhibit mixed results upon external validation in data from ancestries not included in the discovery cohorts

While our clocks demonstrated strong performance on withheld samples from the cohorts that comprised their training sets, we sought to validate their performance in fundamentally distinct data. For this we used samples from the 14 neurologically normal individuals in the Grupo de Neurociencias de Antioquia Brain Bank in Colombia. Ancestry prediction with genotools ^48^, assigned 11 of the 14 as Latino/admixed American ancestry and the remaining 3 as European ancestry. Results are shown in **Supplementary Figure 4**. Among the promoter-based LR-clocks, the NABEC model yielded the strongest correlation (R^2^=0.958), the combined-cohort model produced a similarly strong correlation (R^2^=0.950) while modestly overpredicting chronological age, and the HBCC model resulted in a moderately-strong correlation (R^2^=0.727) with much more drastic age acceleration predictions. The window-based LR-clocks yielded much more stark overpredictions across all three discovery sample sets and negligible correlations. To determine the cause of these near-ubiquitous age overpredictions, we tested the same models on self-normalized validation data, using means and standard deviations from the validation data themselves rather than those which were used for model training. Here we similarly observed strong correlations in the promoter-based models and weak correlations in the window-based models, but with predictions more in line with chronological age, with modest predicted age deceleration across all models. This suggests a handful of promoters/windows with low average methylation in the discovery sample sets may have been included in the models, thereby inflating the normalized values in the validation dataset.

## Discussion

Altogether, the performance of each methylation clock workflow described here demonstrates several key considerations for designing accurate and equitable models. Aggregating methylation across genomic regions, rather than modeling individual CpGs, substantially improved prediction accuracy and cross-cohort generalizability. This can likely be attributed to the reduction of measurement variability through aggregation across neighbouring CpGs. Moreover, the results were much more comparable between the NABEC and HBCC cohorts when using aggregated data than CpG-level data, suggesting that this may have also helped mitigate technical and/or ancestry-specific differences in the methylation datasets. This method was achievable through the enhanced resolution of long-read sequencing data, highlighting its promise in resolving epigenetic complexity. Additionally, models that selected informative regions through data-driven feature selection yielded the most accurate predictions, underscoring the need to account for population-specific epigenetic signatures in biological aging. The overperformance of models utilizing whole-genome windows compared to their analogous promoter-based models in the discovery cohorts underscores the advantages of utilizing features that account for the broader genomic methylation context made possible by long-read sequencing. Additionally, prior clocks developed using loci selected based on brain tissue produced favorable outcomes compared to those whose CpG sets were influenced by non-brain tissue. This is indicative of distinct age-associated methylation patterns across tissues, supporting the value of tissue-specific methylation clocks.

PMI showed no association with age acceleration in any sample set, while cell type composition explained only a modest proportion of variance in age acceleration, with no individual cell type surviving multiple-testing correction. Although both covariates correlate with chronological age, neither substantially predicts deviation from expected epigenetic age. Despite only achieving nominal significance in the HBCC joint model, the consistent negative association between inhibitory neuron fraction and age acceleration across sample sets is notable, as this may reflect the well-documented vulnerability of inhibitory neurons in aging and neurodegeneration ^30^. This suggests that our clocks identify age-related methylation patterns largely independent of sampling biases.

Functional enrichment analyses collectively identify recurrent functional annotations while also showing differences between cohorts. Across each sample set and both feature types, genes mapped to hypermethylated features were enriched for GO terms related to sequence-specific DNA binding and transcriptional regulation, suggesting age-related methylation gains preferentially accumulate at transcriptionally regulated loci. The identification of REST, a master repressor of neuronal gene expression ^49,50^, as the most significantly enriched motif in hypermethylated NABEC and combined-cohort promoters is consistent with prior evidence implicating REST in neuronal aging and neuroprotection. EWCE also identified genes preferentially expressed in astrocytes and OPCs as consistently enriched among hypomethylated and hypermethylated window-based features, respectively, across all sample sets, representing the most generalizable cell type signals identified. EWCE enrichment was substantially broader in NABEC than in HBCC promoter-based models, consistent with GO analyses and likely reflecting the methodological distinction that window-based genes are assigned by direct genomic overlap while promoter-based genes are assigned by annotated associations. Hypomethylated gene sets, though yielding fewer enrichment hits overall, consistently implicated anterior-posterior patterning pathways, with HOX family genes featured prominently. Notably, HOX loci also appeared among hypermethylated regions, consistent with prior reports of bidirectional age-associated methylation changes at HOX clusters ^51^ and suggestive of a broader dysregulation of epigenetic patterning ^52^.

The prominence of PCDH cluster promoters among the strongest predictors in the NABEC and combined-cohort models mirrors prior reports of protocadherin methylation as a hallmark of brain aging, while their near-absence from HBCC models further implicates differences in mechanisms influencing our LR-clocks. The convergence of HOMER, EWCE, and GO findings, particularly in window-based models, around transcriptional regulation, neuronal cell types, and developmentally programmed genomic regions collectively supports a paradigm in which LR-clocks capture the progressive restriction of developmental gene expression programs through both shared and ancestry-specific molecular pathways.

Some limitations warrant consideration. The two discovery cohorts were sequenced using different ONT chemistries (R9 for NABEC; R10 for HBCC), while the Colombian cohort used for external validation was sequenced using a later version of the R10 chemistry, generated in a separate sequencing run. This complicates isolating ancestry as the differentiating variable, since R10 is known to more confidently call methylation at the extrema (0% or 100% methylated) than R9 ^53^, potentially causing systematic miscalibration across chemistries. Consequently, further investigation is required to determine whether differences between the cohort-derived models reflect biological variation, technical effects, differences in cohort composition, or a combination of these factors. Another potential limitation is the difference in age distributions between the cohorts: NABEC exhibited a bimodal distribution, featuring predominantly older (>80) and younger (<20) samples, while HBCC followed a more normal curve almost entirely composed of samples between 20-80 years old. **(Supplementary Figure 13).** This may have enabled the NABEC (and combined-cohort) model to detect age-predictive features associated with early development and advanced age that were less readily detectable in HBCC-specific models, including the absence of the REST motif in HBCC-specific models. Despite promoter-based models producing predictions that strongly correlated with chronological age, the HBCC and, to a lesser degree, combined-cohort models consistently overpredicted age. This may reflect technical artifacts uniformly impacting HBCC samples while still preserving the majority of age-associated variation in methylation patterns. Although window-based LR-clocks performed better within their training cohorts, they failed to generalize to new cohorts as effectively as promoter-based models. This disparity suggests promoter-level methylation patterns may be more conserved across populations, undergoing more predictable increases with age. Conversely, non-promoter loci across the genome appear to exhibit greater epigenetic drift that may be unique to specific populations. These results underscore the necessity of including diverse samples in training datasets, particularly when using annotation-agnostic genome-wide features. In addition, the biorepositories used do not consistently capture real-world clinical information such as longitudinal or electronic medical record data, and cause of death and comorbid illness burden are only sparsely quantified, which may contribute to sample heterogeneity. As long-read methylation datasets grow, future clocks will benefit from larger sample sizes, harmonized sequencing protocols, and integration with comprehensive phenotypic data.

Together, these findings demonstrate that long-read methylation profiling enables accurate, ancestry-aware modeling of biological aging in the brain. Our results reinforce the need for inclusive, population-representative datasets and suggest novel cell type–specific pathways involved in aging. Finally, these workflows were developed in accordance with open science principles, with code publicly available at https://zenodo.org/records/21777087 and https://github.com/NIH-CARD/LongRead-Methylation-Clocks.

## Methods

### Discovery cohort long-read DNA methylation data generation

We leveraged existing ONT WGS data generated from two postmortem brain cohorts: the North American Brain Expression Consortium (NABEC) and the Human Brain Collection Core (HBCC). HBCC samples were collected at the National Institute of Mental Health Intramural Research Program, while NABEC samples were collected across the United States at the Banner Sun Health Research Institute, the University of Kentucky Alzheimer’s Disease Center Brain Bank, and the University of Maryland Brain and Tissue bank. Full details of the experimental protocol are available elsewhere ^54^. In brief, genomic DNA was extracted from PFC tissue dissected from frozen brain samples for 130 neurologically healthy individuals from HBCC and 187 neurologically healthy individuals from NABEC ^54^. ONT PromethION sequencing was performed on all samples using R9.4.1 flow cells basecalled with Guppy v6.12 for NABEC and R10.4.1 flow cells basecalled with Guppy v6.38 for HBCC; both cohorts included 5mC methylation in the basecalling command. Samples were sequenced until they reached at least 115 GB of sequencing data. All samples were processed through the Nanopore Analysis Pipeline (Napu) ^55^, wherein reads were aligned to GRCh38 with minimap2 ^56^ and methylation BED files were then produced using Modkit (https://github.com/nanoporetech/modkit) pileup to CpG motifs in the same reference. Methylation frequency at each CpG site was defined as the proportion of aligned reads carrying a methylated cytosine at that position. Genetic ancestry was determined using genotools ancestry prediction ^48^.

### External validation cohort long-read DNA methylation data generation

In the Colombian cohort, from the Grupo de Neurociencias de Antioquia Brain Bank, ONT PromethION sequencing was performed on middle frontal gyrus tissue from 14 neurologically healthy individuals, ranging from 38 to 90+ years old. For sequencing, R10.4.1 flowcells were used and 30x read coverage was targeted. For basecalling with 5mC methylation modification calls, dorado v0.5.0 (model: dna_r10.4.1_e8.2_400bps_sup@v4.2.0 5mCG_5hmCG) was used. Reads were aligned to GRCh38 with minimap2 ^56^ and methylation BED files were generated using Modkit (https://github.com/nanoporetech/modkit) pileup to CpG motifs in the same reference. Methylation frequency at each CpG site was defined as the proportion of aligned reads carrying a methylated cytosine at that position. The project was approved by the Institutional Review Board (IRB) of the Medical Research institute, School of Medicine, Universidad de Antioquia. Written informed consent following the guidelines of the Code of Ethics of the World Medical Association, Helsinki declaration, and Belmont Report was obtained from all participants or their legally authorized proxies.

### Methylation data processing and feature aggregation

CpG sites were filtered within each cohort to only include those with a coverage minimum of 3 reads across both haplotypes for at least 95% of the participants in the respective cohort. To ensure that we use an equal number of CpGs in all samples, CpGs within deletions were removed from all samples. To do this deletions were isolated from a merged set of cohort SVs, converted into a bed file, and used to remove CpGs within cohort deletions from the cohort methylation data prior to aggregating. Promoter annotations were obtained from the Eukaryotic Promoter Database (EPD) ^28^. These 60-basepair EPDnew promoter regions were expanded by 1 kb on each end, yielding regions centered on transcription start sites to encompass a broader CpG context. EPDnew promoter identifiers comprise the associated gene symbol followed by a numerical suffix representing the promoter-usage hierarchy for that gene, with “_1” denoting the highest-usage promoter. Whole-genome windows were created by dividing chromosomes into non-overlapping windows containing at least 50 CpGs, starting with a minimum width of 1 kb, as has been described elsewhere ^54^. Windows initially containing less than 50 CpGs were extended until 50 CpGs were included. Methylation was then aggregated using a weighted mean across CpG sites within each of these promoter regions ^57^, yielding data for 27,620; 27,524; and 27,285 promoters and 502,300; 502,007; and 494,831 whole-genome windows in the HBCC, NABEC, and combined-cohort sample sets, respectively. Aggregating methylation frequencies across genomic regions reduces noise from individual CpG measurements, enhances statistical power, and enables interpretation of methylation at the level of regulatory elements.

In each of the promoter and whole-genome window models, features were then filtered according to their correlation with other features and their independent associations with age using the munging module in GenoML v2.0.0. For each model, age was regressed against each feature individually using ordinary least squares (OLS) to determine which are most likely to be linked with aging. Only those with significant (p < 0.05) predictive power were kept. Pearson’s correlation was then used to find correlation values for each pair of remaining features. Pairs with high correlation values (r > 0.9) were deemed to be redundant, and thus only the feature with the lower OLS p-value of each respective pair was kept. These filtered and aggregated methylation features were subsequently used as inputs for machine learning model development.

### Comparisons with existing clocks

To evaluate the performance of existing methylation clocks using long-read methylation data, LR-clocks were benchmarked against several previously-reported methylation clocks, including the Horvath pan-tissue clock ^1^, PhenoAge ^3^, DunedinPACE ^5^, the three pan-mammalian pan-tissue clocks from Lu *et al.* ^2^, and the brain-specific clock from Shireby *et al.* ^24^. To compare our methodology, the CpG sites that were used to develop each respective methylation clock were extracted from the long-read datasets used in this study. For any CpG sites from these clocks which were not captured in the datasets used here, the nearest CpG site was used in its place. New models were developed according to the workflow described here using those individual CpG sites, as well as finding the corresponding whole-genome window and the nearest promoter to each of those loci, and using those as inputs. For instances in which two CpG sites from the same benchmark model shared the same whole-genome window or nearest promoter, each feature was only included once in the input dataset for each respective model.

### Principal component models

To develop PCA-based models, we performed PCA on the long-read dataset for each sample set, with methylation fraction at each CpG which met QC criteria for a given sample set. This amounted to more than 22 million CpGs included for each dataset, which is both prohibitively large for PCA and likely includes low-variance CpGs that would not meaningfully contribute to the PCA. Therefore, we kept only the CpGs in the top quartile of variance in each dataset and performed PCA. We defined the number of principal components for each dataset as being equal to the number of samples, as has been discussed elsewhere ^14^.

### Statistical analysis and machine learning

For each clock model, the respective methylation data were used as inputs for machine learning model development using GenoML v2.0.0, which is described elsewhere ^27^. In brief, GenoML consists of separate modules for munging (which handles data preprocessing), training (which competes several model architectures against each other to identify the best-performing model class), tuning (which scans the hyperparameter space to find those which combination of hyperparameters best fits each respective dataset), and testing (which applies the final model to withheld data). For this study, all analyses were run with a random state of 3 and with nested cross-validation (CV), using 5 folds for both the inner and outer CV steps. This is implemented in GenoML by defining the outer CV split in the munging step, stratified by phenotype (using decile bins, as age is a continuous variable) and, for the LR-clocks, using values of 0.05 and 0.9 for OLS p-value and Pearson’s r coefficient thresholds, respectively, for feature filtering. Munging is performed on each fold, as well as the full dataset, separately to prevent data leakage. Then, we performed a 70:30 train:validation split on the full dataset (to ensure the same architecture is used across all models from the same dataset) during the training phase to determine the model class to be used. This was followed by model tuning, wherein for each of the five outer CV folds (plus the full dataset) another 5-fold inner CV was performed for hyperparameter optimization, yielding six different models with different combinations of hyperparameters, each corresponding to a different split (or lack thereof) of the full dataset. Tuning was performed with 50 candidate hyperparameter iterations using BayesSearchCV. The testing module was then used to evaluate the performance of the withheld data from each outer CV fold, with the model developed on the full dataset being used for deployment as well as for downstream feature importance and functional enrichment analyses.

### Feature importance modeling

Feature importance quantifications for each of the LR-clocks were ascertained using the SHapley Additive exPlanations (SHAP) package v0.48.0 ^58^. SHAP utilizes a game theoretic model to assign a “SHAP value” to each feature, representing how much it contributes to the final prediction of the model. These were visualized as beeswarm plots, which depict sample-level SHAP values color-coded to indicate directionality of effect, as well as barplots, which show the average magnitude of effect combined across all samples to indicate the features with the strongest overall impact. All plots include the 20 features with the highest average SHAP value magnitude for each model.

### Gene set Enrichment Analysis

Genes of interest derived from each of the LR-clocks were functionally annotated with gene ontology (GO) terms using the gProfiler2 R (v4.3) package. Background gene lists consisted of genes included as inputs into models after filtering features based on age association and correlation with other features. Terms consisting of fewer than 10 or more than 2000 genes were removed, and p-values were corrected for multiple testing using the recommended g:SCS method. GO terms were reduced to parent terms using semantic similarity with a cut-off of 0.5, using the Rutils GitHub package (https://github.com/RHReynolds/rutils) to aid visualisation.

### Expression weighted cell type enrichment

We performed cell-type enrichment using Expression Weighted Cell Type Enrichment (EWCE, v. 1.4.0) ^59^ to assess whether our genes of interest show significantly higher expression in specific cell types than expected by chance. This analysis was conducted on single-nucleus transcriptomes from 54,394 nuclei from the human dorsolateral prefrontal cortex (DLPFC) without neurological disorders. Preprocessing and annotation of this dataset is reported elsewhere ^60^, and the function generate_celltype_data() from the R (4.2.0) package EWCE was used to generate the celltype dataset.

Bootstrap gene lists controlled for transcript length and GC-content were generated with EWCE iteratively (n=10,000) using the “bootstrap_enrichment_test()” function. This function takes the inquiry gene list and a single cell type transcriptome data set and determines the probability of enrichment of this list in a given cell type when compared to the gene expression of bootstrapped gene lists, returning the probability of enrichment and fold-change of enrichment. P-values were corrected for multiple testing using the false discovery rate (fdr) method.

### Transcription factor binding motif enrichment

To identify the TF-binding motifs with the strongest influence in each of our LR-clocks, we used HOMER v4.11 ^61^. Our promoters of interest were defined as any promoter (or promoters overlapping with any whole-genome window) that yielded a nonzero SHAP value for each respective model. Background promoters were defined as all promoters (or promoters overlapping with any whole-genome window) that passed QC for each respective sample set.

### Covariate regression analysis

Age acceleration for each participant was computed as the residual from an OLS regression of predicted age on chronological age, fitted separately within each cohort. This was done to remove systematic bias in clock predictions across the age distribution and isolate individual-level deviation from expected epigenetic age. Age acceleration was regressed against PMI and estimated composition of seven canonical cell types (Astrocytes, Excitatory Neurons, Inhibitory Neurons, Microglia, OPCs, Oligodendrocytes, and Vascular Cells), which were determined as described elsewhere ^29^ for the 129 HBCC and 167 NABEC samples with single-cell transcriptomic data. Individual models were fit for each covariate separately, and a joint model including all cell type fractions simultaneously was fit with one cell type omitted as a reference to avoid perfect collinearity. A further joint model including both PMI and all cell type fractions was fit to assess independent effects. P-values were corrected for multiple testing across all variables using the Benjamini-Hochberg false discovery rate method.

## Data availability

Human brain sequencing datasets are under controlled access and require a dbGap application (phs001300.v5) (phs000979.v4). Afterwards, the data will be available through the restricted AnVIL workspace.

## Acknowledgements

This research was supported [in part] by the Intramural Research Program of the National Institutes of Health (NIH). The contributions of the NIH author(s) were made as part of their official duties as NIH federal employees, are in compliance with agency policy requirements, and are considered Works of the United States Government. However, the findings and conclusions presented in this paper are those of the author(s) and do not necessarily reflect the views of the NIH or the U.S. Department of Health and Human Services. The HBCC is funded by the NIMH-IRP through project ZIC MH002903. This work utilized the computational resources of the NIH HPC Biowulf cluster (https://hpc.nih.gov). We thank members of the North American Brain Expression Consortium (NABEC) for providing samples derived from brain tissue. We are grateful to the Banner Sun Health Research Institute Brain and Body Donation Program of Sun City, Arizona for the provision of human biological materials. The Brain and Body Donation Program has been supported by the National Institute of Neurological Disorders and Stroke (U24 NS072026 National Brain and Tissue Resource for Parkinson’s Disease and Related Disorders), the National Institute on Aging (P30 AG19610 and P30AG072980, Arizona Alzheimer’s Disease Center), University of Kentucky Alzheimer’s Disease Center Brain Bank NIA P30 AG072946, the Arizona Department of Health Services (contract 211002, Arizona Alzheimer’s Research Center), the Arizona Biomedical Research Commission (contracts 4001, 0011, 05-901 and 1001 to the Arizona Parkinson’s Disease Consortium). We thank the Biobank of the Grupo de Neurosciencias de Antioquia, which has been supported by the U.S. National Institute on Aging (R01AG062479). KJB has been in part supported by William H. Gates Sr. Fellowships from the Alzheimer’s Disease Data Initiative. S.J.E. is supported by American Heart Association Grant #26PRE1545631 / Sarah Eger / 2026.

## Author contributions

M.J., L.F., M.R.C., M.A.N., and K.J.B. conceptualized the study. S.M.G., M.B.M., M.M., H.L., N.K., C.W., X.R., M.J., L.F., M.R.C., M.A.N., and K.J.B. designed the study. M.M., Ab.M., Aj.M., and P.A., A.V-L., B.M-C., J.A-U., D.A., and K.S.K. contributed to data acquisition. S.M.G., S.J.E., M.B.M., M.M., Ab.M., X.R., and A.C. contributed to data preprocessing. S.M.G., S.J.E., M.B.M., M.G.P., and A.H. performed the bioinformatics analyses. L.F., M.R., M.A.N., K.J.B. provided critical input on the results. S.M.G., M.B.M., M.M., M.A.N., and K.J.B. drafted the paper. All authors reviewed and approved the final version.

