## Supplementary Figures for "Long-Read epigenetic clocks identify improved brain aging predictions"

### Slide 1
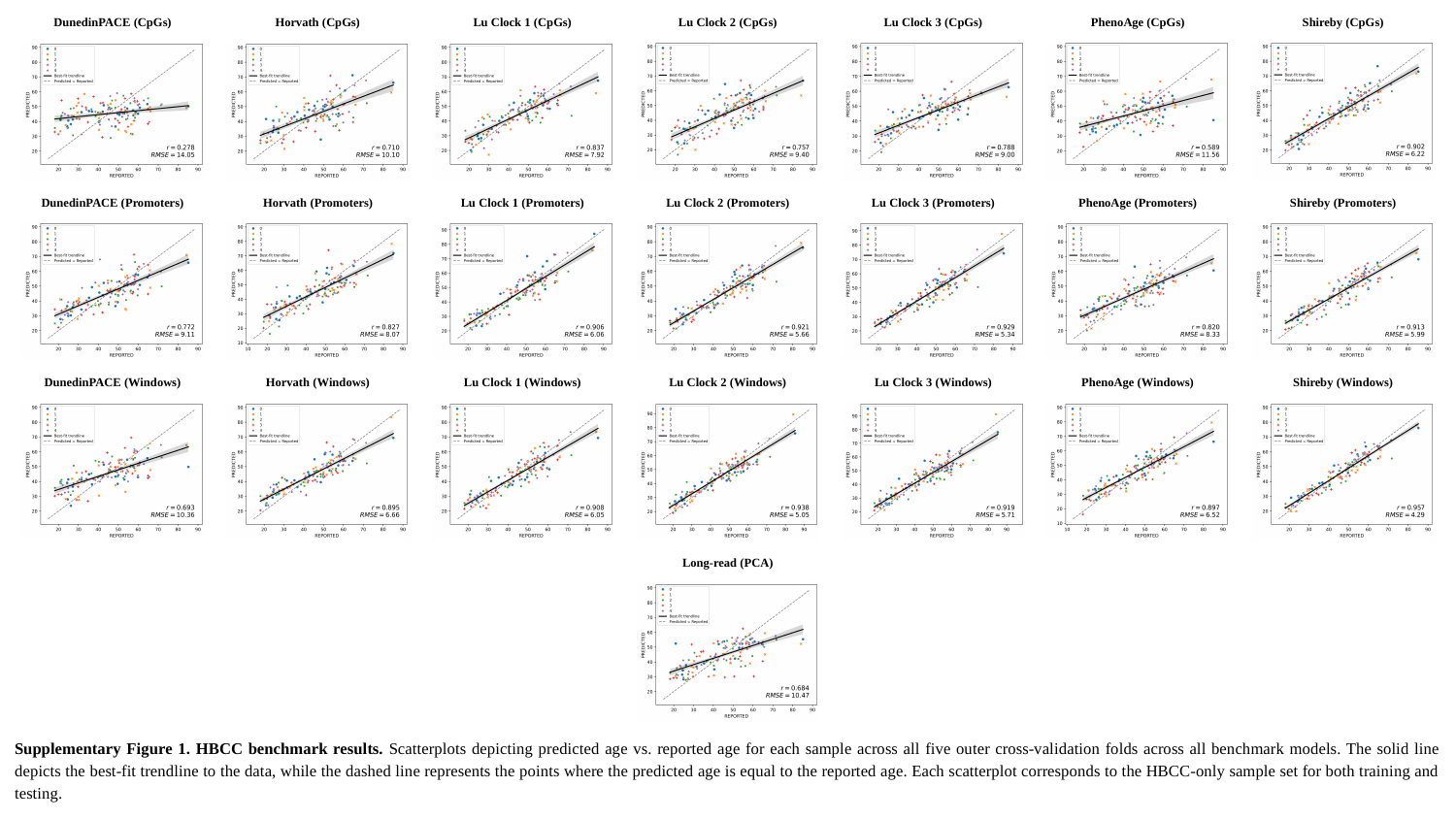

Shireby (CpGs)
Lu Clock 3 (CpGs)
Lu Clock 2 (CpGs)
DunedinPACE (CpGs)
Horvath (CpGs)
Lu Clock 1 (CpGs)
PhenoAge (CpGs)
Lu Clock 3 (Promoters)
Shireby (Promoters)
Lu Clock 2 (Promoters)
DunedinPACE (Promoters)
Horvath (Promoters)
Lu Clock 1 (Promoters)
PhenoAge (Promoters)
Lu Clock 3 (Windows)
Shireby (Windows)
Lu Clock 2 (Windows)
DunedinPACE (Windows)
Horvath (Windows)
Lu Clock 1 (Windows)
PhenoAge (Windows)
Long-read (PCA)
Supplementary Figure 1. HBCC benchmark results. Scatterplots depicting predicted age vs. reported age for each sample across all five outer cross-validation folds across all benchmark models. The solid line depicts the best-fit trendline to the data, while the dashed line represents the points where the predicted age is equal to the reported age. Each scatterplot corresponds to the HBCC-only sample set for both training and testing.

### Slide 2
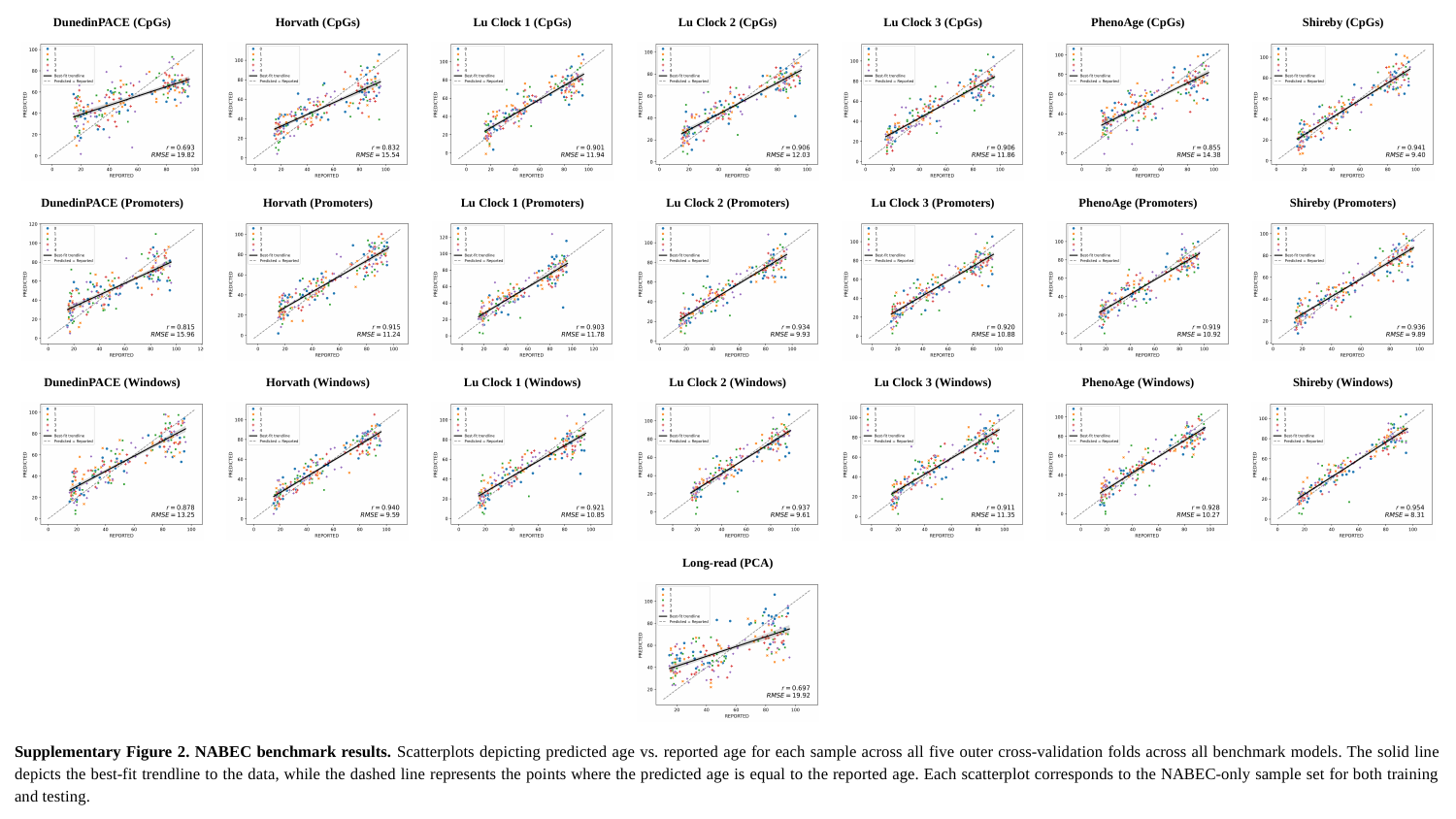

DunedinPACE (CpGs)
Horvath (CpGs)
Lu Clock 1 (CpGs)
Lu Clock 2 (CpGs)
Lu Clock 3 (CpGs)
Shireby (CpGs)
PhenoAge (CpGs)
DunedinPACE (Promoters)
Horvath (Promoters)
Lu Clock 1 (Promoters)
Lu Clock 2 (Promoters)
Lu Clock 3 (Promoters)
Shireby (Promoters)
PhenoAge (Promoters)
DunedinPACE (Windows)
Horvath (Windows)
Lu Clock 1 (Windows)
Lu Clock 2 (Windows)
Lu Clock 3 (Windows)
Shireby (Windows)
PhenoAge (Windows)
Long-read (PCA)
Supplementary Figure 2. NABEC benchmark results. Scatterplots depicting predicted age vs. reported age for each sample across all five outer cross-validation folds across all benchmark models. The solid line depicts the best-fit trendline to the data, while the dashed line represents the points where the predicted age is equal to the reported age. Each scatterplot corresponds to the NABEC-only sample set for both training and testing.

### Slide 3
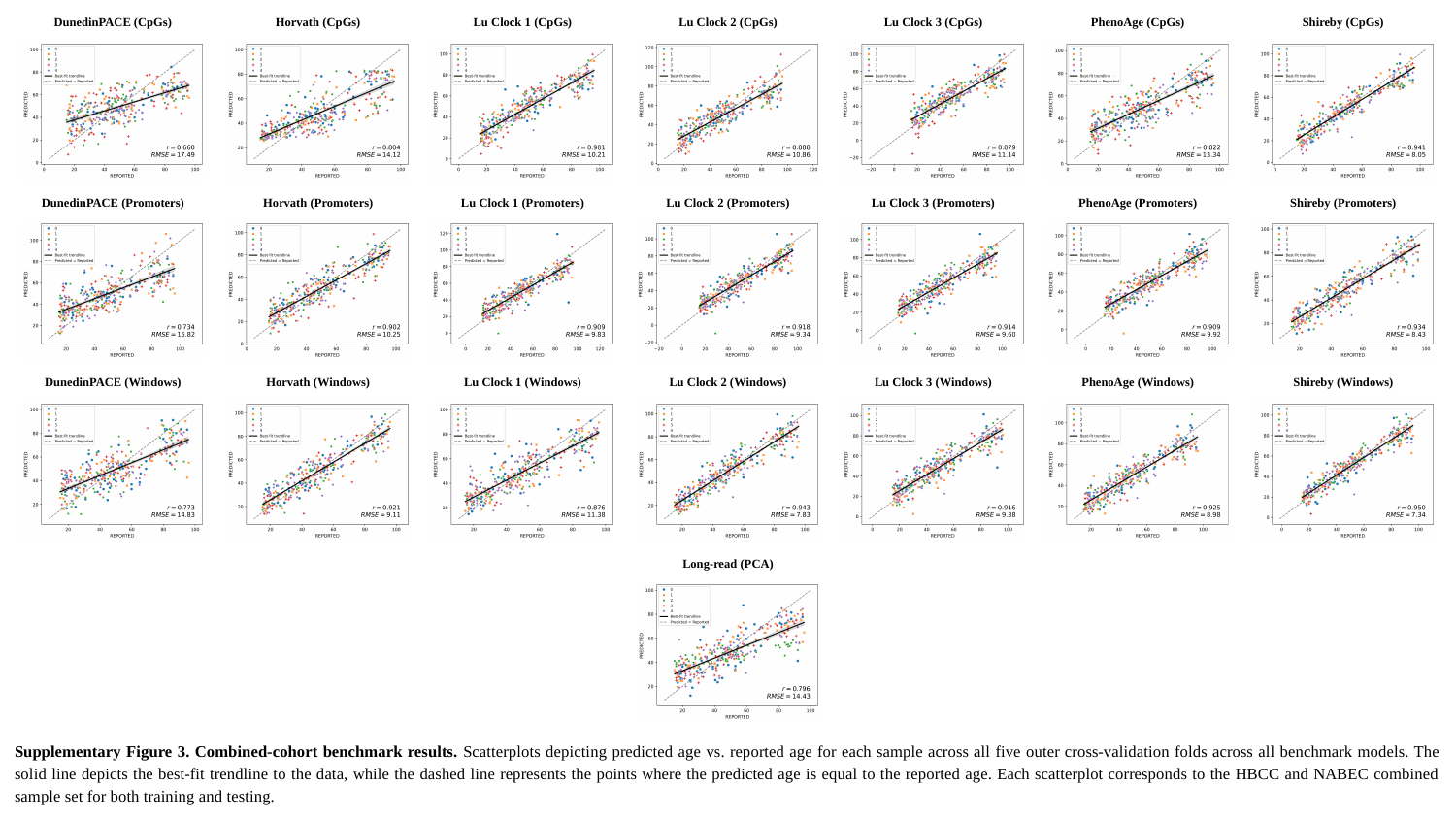

Shireby (CpGs)
DunedinPACE (CpGs)
Horvath (CpGs)
Lu Clock 1 (CpGs)
Lu Clock 2 (CpGs)
Lu Clock 3 (CpGs)
PhenoAge (CpGs)
DunedinPACE (Promoters)
Horvath (Promoters)
Lu Clock 1 (Promoters)
Lu Clock 2 (Promoters)
Lu Clock 3 (Promoters)
Shireby (Promoters)
PhenoAge (Promoters)
DunedinPACE (Windows)
Horvath (Windows)
Lu Clock 1 (Windows)
Lu Clock 2 (Windows)
Lu Clock 3 (Windows)
Shireby (Windows)
PhenoAge (Windows)
Long-read (PCA)
Supplementary Figure 3. Combined-cohort benchmark results. Scatterplots depicting predicted age vs. reported age for each sample across all five outer cross-validation folds across all benchmark models. The solid line depicts the best-fit trendline to the data, while the dashed line represents the points where the predicted age is equal to the reported age. Each scatterplot corresponds to the HBCC and NABEC combined sample set for both training and testing.

### Slide 4
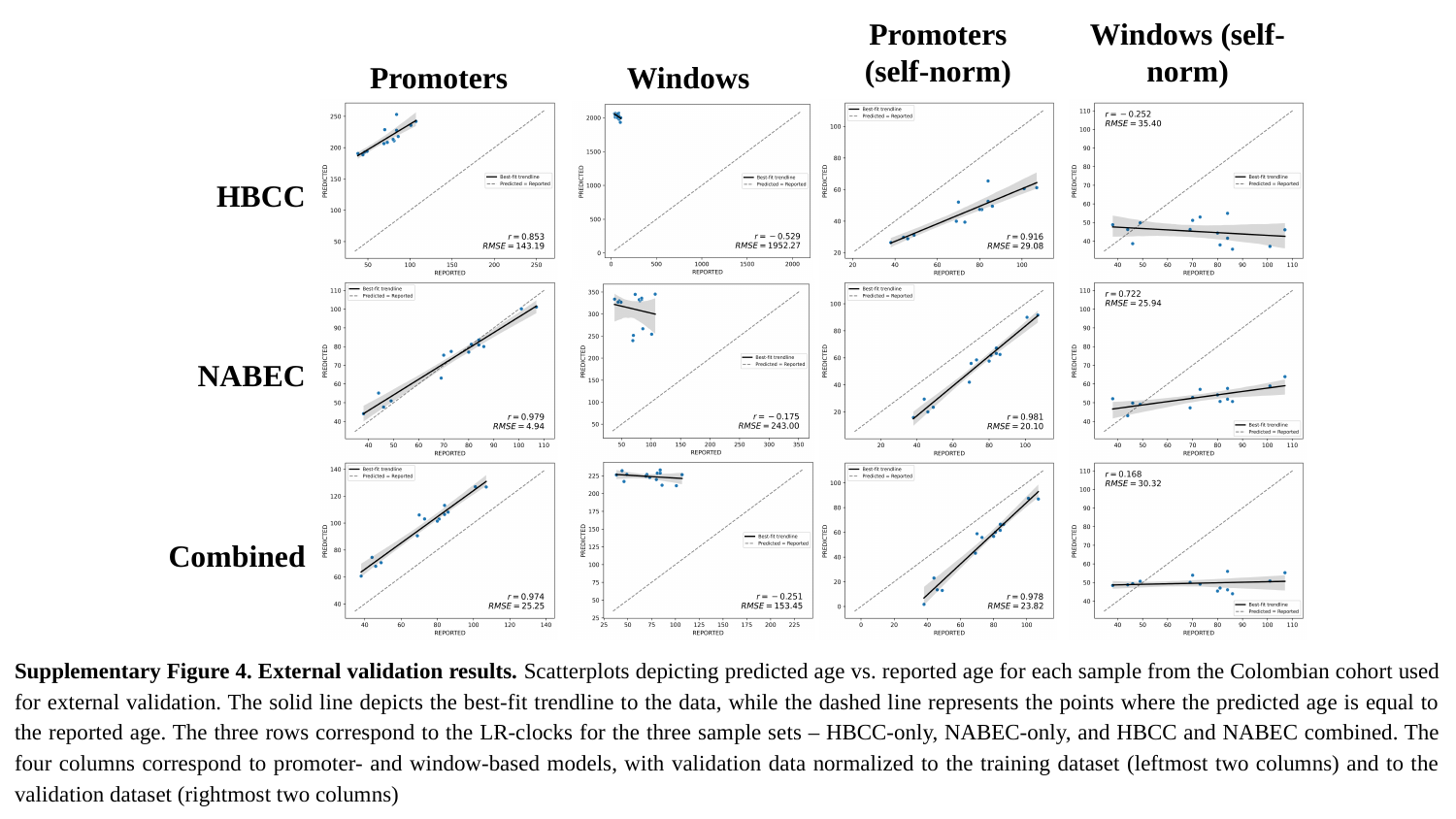

Promoters (self-norm)
Windows (self-norm)
Promoters
Windows
HBCC
NABEC
Combined
Supplementary Figure 4. External validation results. Scatterplots depicting predicted age vs. reported age for each sample from the Colombian cohort used for external validation. The solid line depicts the best-fit trendline to the data, while the dashed line represents the points where the predicted age is equal to the reported age. The three rows correspond to the LR-clocks for the three sample sets – HBCC-only, NABEC-only, and HBCC and NABEC combined. The four columns correspond to promoter- and window-based models, with validation data normalized to the training dataset (leftmost two columns) and to the validation dataset (rightmost two columns)

### Slide 5
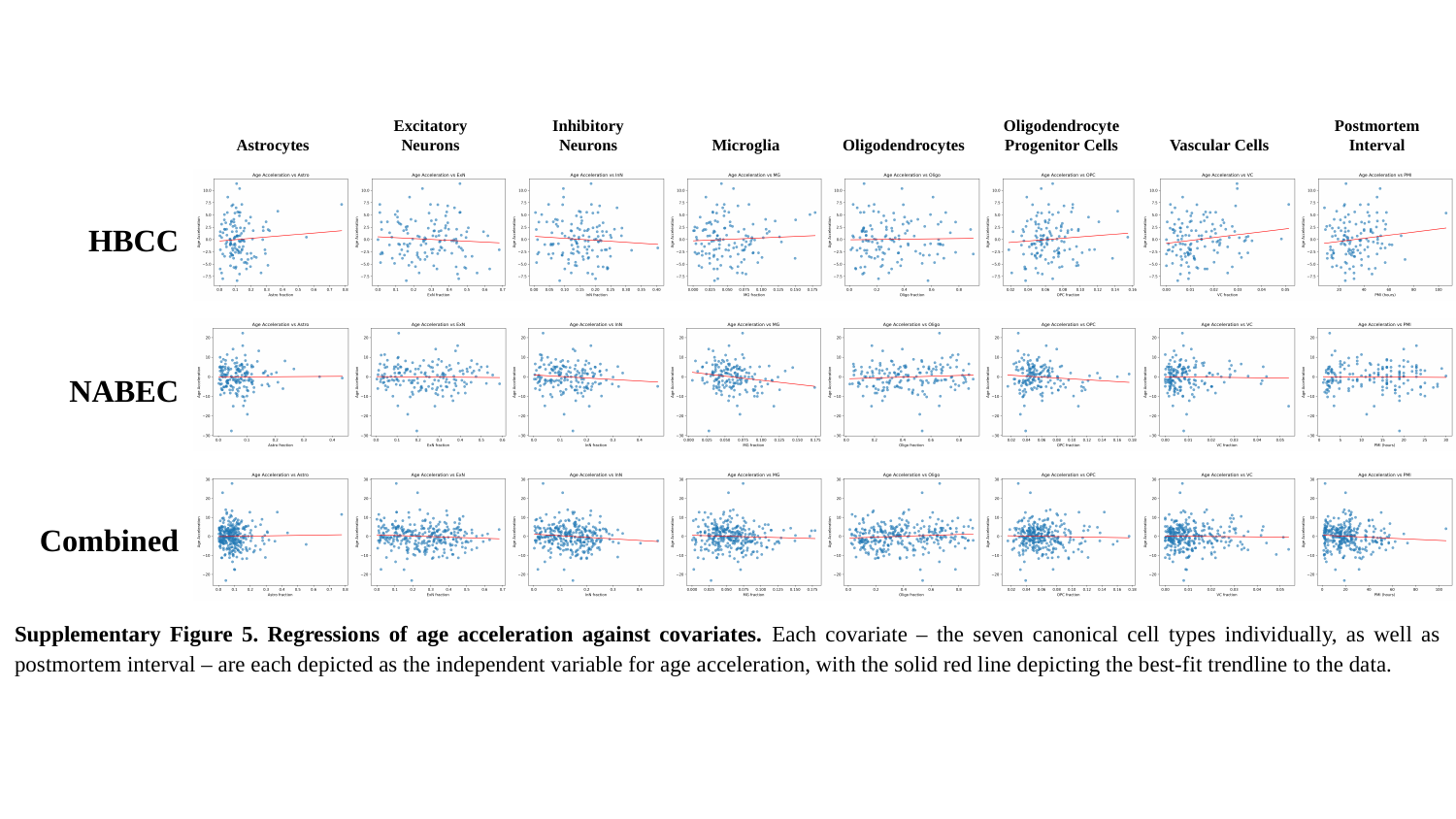

Astrocytes
Excitatory Neurons
Inhibitory Neurons
Microglia
Oligodendrocytes
Oligodendrocyte Progenitor Cells
Vascular Cells
Postmortem Interval
HBCC
NABEC
Combined
Supplementary Figure 5. Regressions of age acceleration against covariates. Each covariate – the seven canonical cell types individually, as well as postmortem interval – are each depicted as the independent variable for age acceleration, with the solid red line depicting the best-fit trendline to the data.

### Slide 6
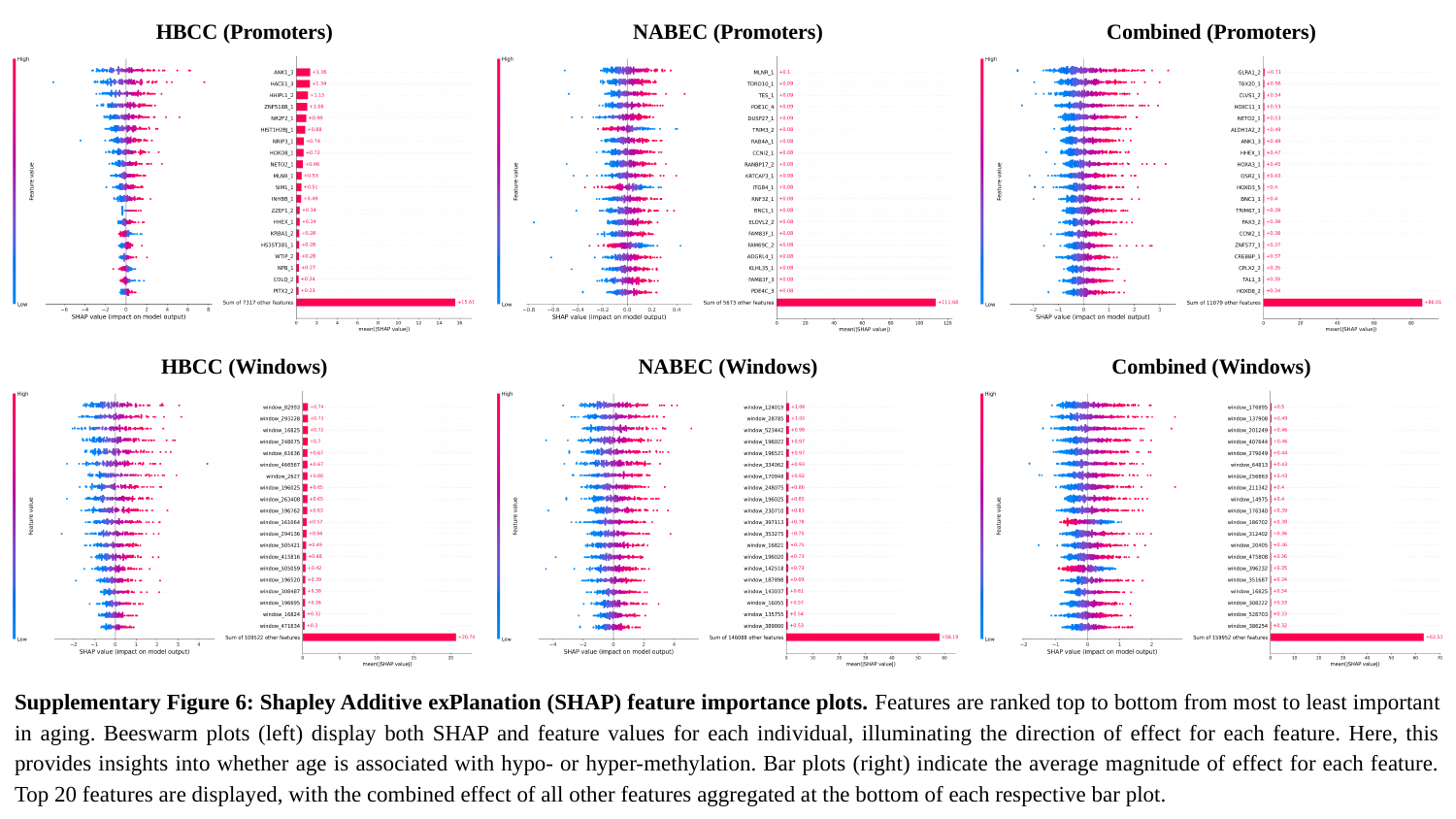

HBCC (Promoters)
NABEC (Promoters)
Combined (Promoters)
HBCC (Windows)
NABEC (Windows)
Combined (Windows)
Supplementary Figure 6: Shapley Additive exPlanation (SHAP) feature importance plots. Features are ranked top to bottom from most to least important in aging. Beeswarm plots (left) display both SHAP and feature values for each individual, illuminating the direction of effect for each feature. Here, this provides insights into whether age is associated with hypo- or hyper-methylation. Bar plots (right) indicate the average magnitude of effect for each feature. Top 20 features are displayed, with the combined effect of all other features aggregated at the bottom of each respective bar plot.

### Slide 7
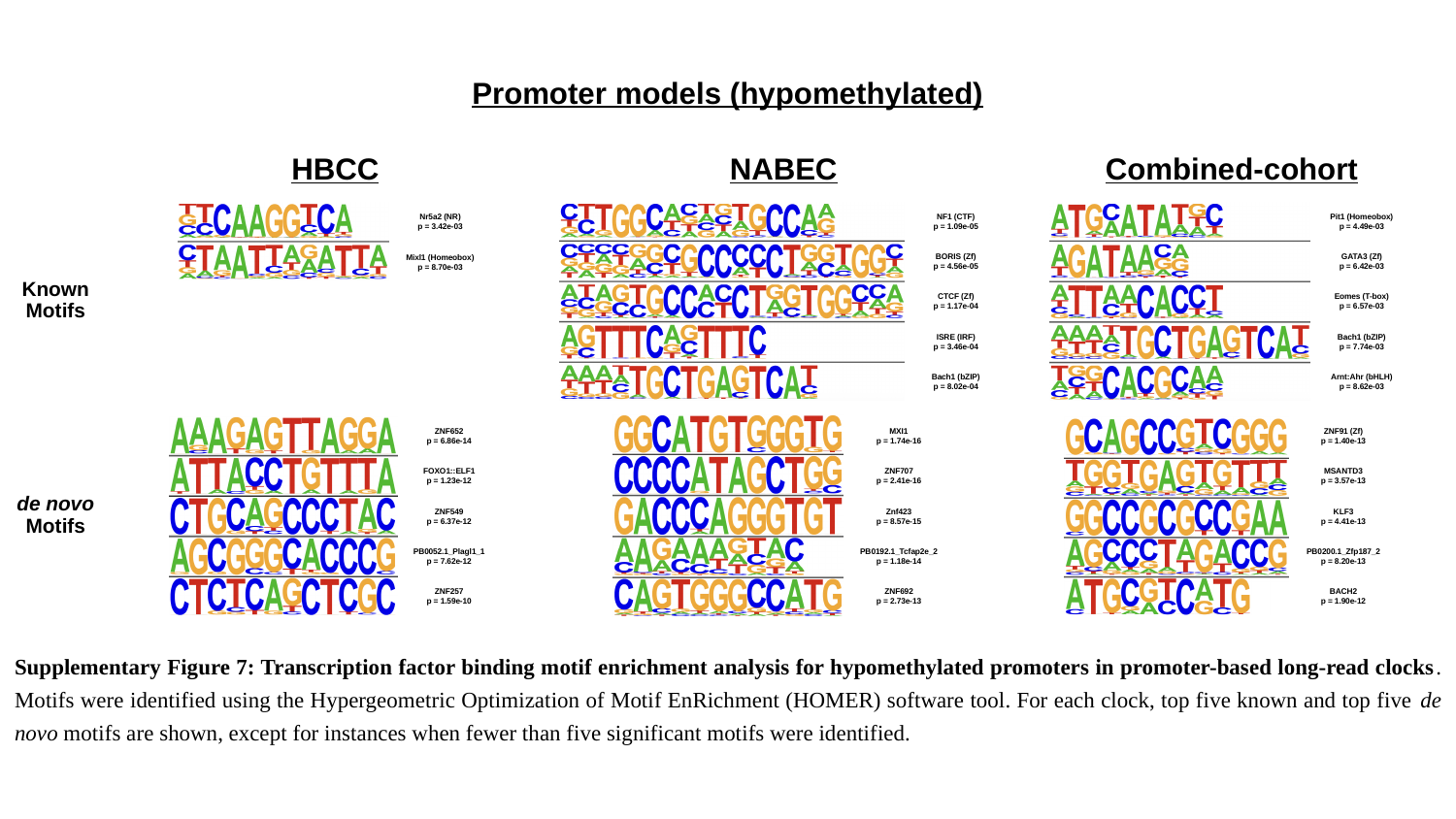

Promoter models (hypomethylated)
HBCC
NABEC
Combined-cohort
NF1 (CTF)
p = 1.09e-05
BORIS (Zf)
p = 4.56e-05
CTCF (Zf)
p = 1.17e-04
ISRE (IRF)
p = 3.46e-04
Bach1 (bZIP)
p = 8.02e-04
Pit1 (Homeobox)
p = 4.49e-03
GATA3 (Zf)
p = 6.42e-03
Eomes (T-box)
p = 6.57e-03
Bach1 (bZIP)
p = 7.74e-03
Arnt:Ahr (bHLH)
p = 8.62e-03
Nr5a2 (NR)
p = 3.42e-03
Mixl1 (Homeobox)
p = 8.70e-03
Known Motifs
MXI1
p = 1.74e-16
ZNF707
p = 2.41e-16
Znf423
p = 8.57e-15
PB0192.1_Tcfap2e_2
p = 1.18e-14
ZNF692
p = 2.73e-13
ZNF652
p = 6.86e-14
FOXO1::ELF1
p = 1.23e-12
ZNF549
p = 6.37e-12
PB0052.1_Plagl1_1
p = 7.62e-12
ZNF257
p = 1.59e-10
ZNF91 (Zf)
p = 1.40e-13
MSANTD3
p = 3.57e-13
KLF3
p = 4.41e-13
PB0200.1_Zfp187_2
p = 8.20e-13
BACH2
p = 1.90e-12
de novo Motifs
Supplementary Figure 7: Transcription factor binding motif enrichment analysis for hypomethylated promoters in promoter-based long-read clocks. Motifs were identified using the Hypergeometric Optimization of Motif EnRichment (HOMER) software tool. For each clock, top five known and top five de novo motifs are shown, except for instances when fewer than five significant motifs were identified.

### Slide 8
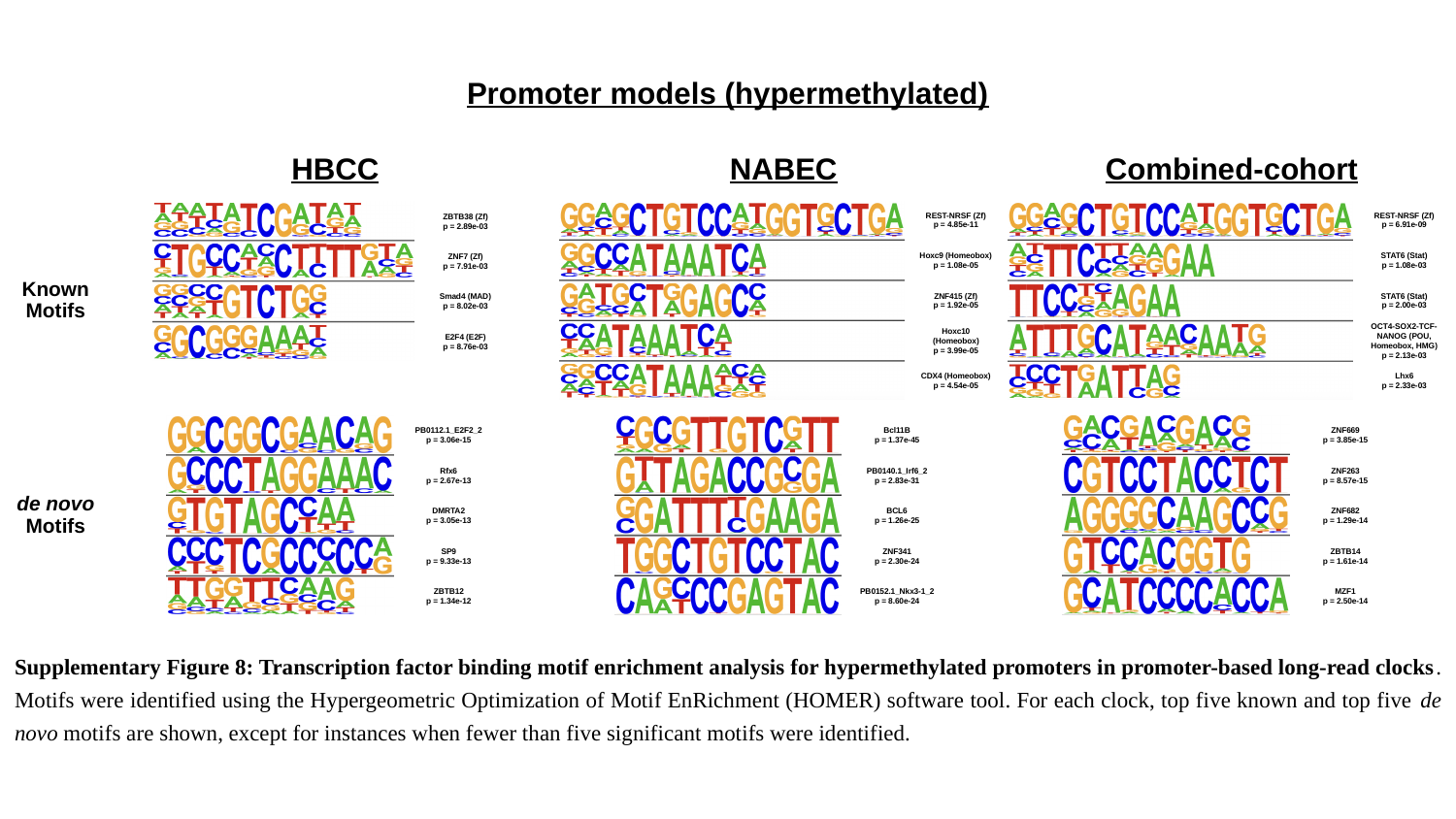

Promoter models (hypermethylated)
HBCC
NABEC
Combined-cohort
REST-NRSF (Zf)
p = 4.85e-11
Hoxc9 (Homeobox)
p = 1.08e-05
ZNF415 (Zf)
p = 1.92e-05
Hoxc10 (Homeobox)
p = 3.99e-05
CDX4 (Homeobox)
p = 4.54e-05
REST-NRSF (Zf)
p = 6.91e-09
STAT6 (Stat)
p = 1.08e-03
STAT6 (Stat)
p = 2.00e-03
OCT4-SOX2-TCF-NANOG (POU, Homeobox, HMG)
p = 2.13e-03
Lhx6
p = 2.33e-03
ZBTB38 (Zf)
p = 2.89e-03
ZNF7 (Zf)
p = 7.91e-03
Smad4 (MAD)
p = 8.02e-03
E2F4 (E2F)
p = 8.76e-03
Known Motifs
PB0112.1_E2F2_2
p = 3.06e-15
Rfx6
p = 2.67e-13
DMRTA2
p = 3.05e-13
SP9
p = 9.33e-13
ZBTB12
p = 1.34e-12
ZNF669
p = 3.85e-15
ZNF263
p = 8.57e-15
ZNF682
p = 1.29e-14
ZBTB14
p = 1.61e-14
MZF1
p = 2.50e-14
Bcl11B
p = 1.37e-45
PB0140.1_Irf6_2
p = 2.83e-31
BCL6
p = 1.26e-25
ZNF341
p = 2.30e-24
PB0152.1_Nkx3-1_2
p = 8.60e-24
de novo Motifs
Supplementary Figure 8: Transcription factor binding motif enrichment analysis for hypermethylated promoters in promoter-based long-read clocks. Motifs were identified using the Hypergeometric Optimization of Motif EnRichment (HOMER) software tool. For each clock, top five known and top five de novo motifs are shown, except for instances when fewer than five significant motifs were identified.

### Slide 9
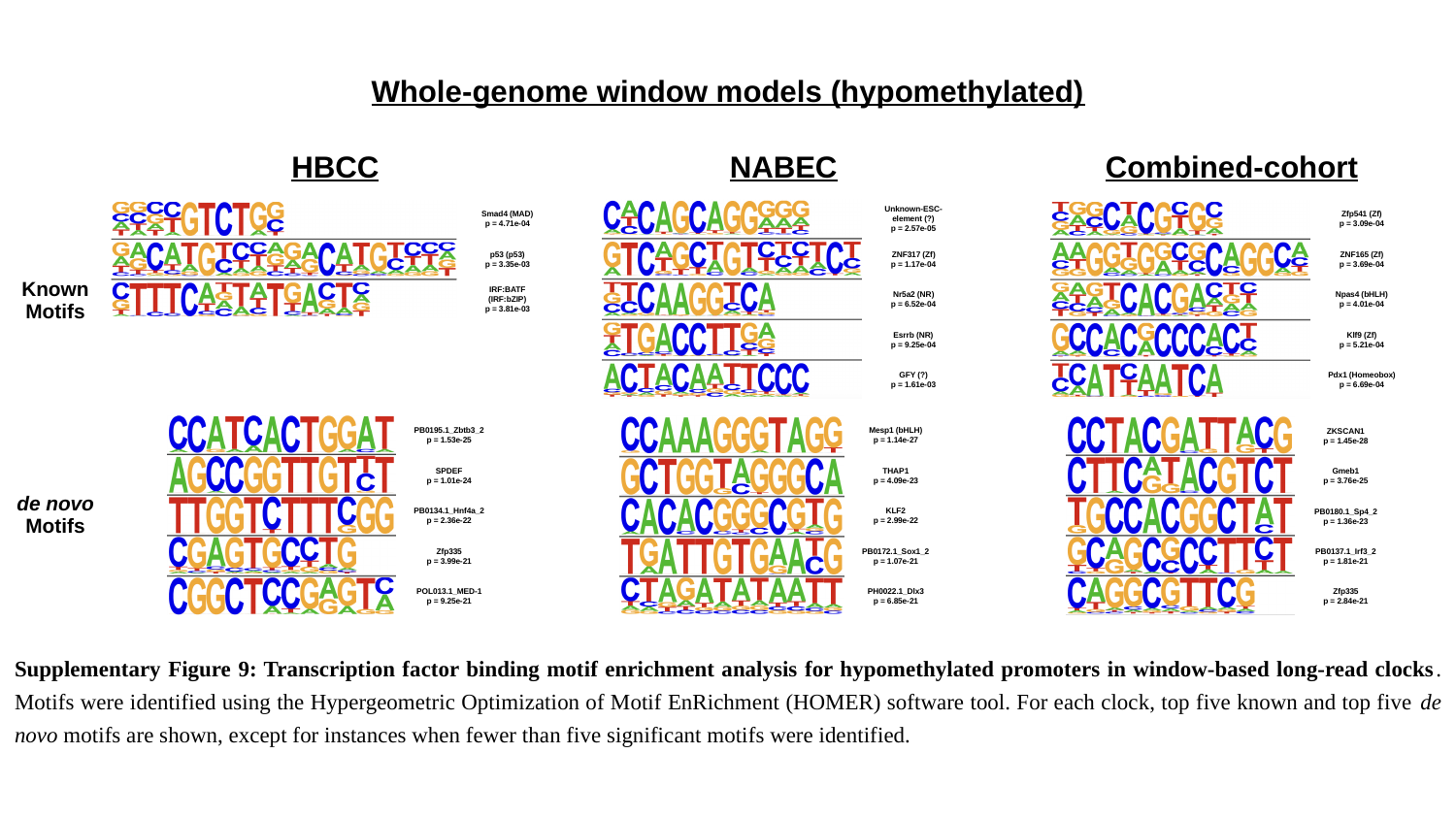

Whole-genome window models (hypomethylated)
HBCC
NABEC
Combined-cohort
Unknown-ESC-
element (?)
p = 2.57e-05
ZNF317 (Zf)
p = 1.17e-04
Nr5a2 (NR)
p = 6.52e-04
Esrrb (NR)
p = 9.25e-04
GFY (?)
p = 1.61e-03
Zfp541 (Zf)
p = 3.09e-04
ZNF165 (Zf)
p = 3.69e-04
Npas4 (bHLH)
p = 4.01e-04
Klf9 (Zf)
p = 5.21e-04
Pdx1 (Homeobox)
p = 6.69e-04
Smad4 (MAD)
p = 4.71e-04
p53 (p53)
p = 3.35e-03
IRF:BATF (IRF:bZIP)
p = 3.81e-03
Known Motifs
PB0195.1_Zbtb3_2
p = 1.53e-25
SPDEF
p = 1.01e-24
PB0134.1_Hnf4a_2
p = 2.36e-22
Zfp335
p = 3.99e-21
POL013.1_MED-1
p = 9.25e-21
Mesp1 (bHLH)
p = 1.14e-27
THAP1
p = 4.09e-23
KLF2
p = 2.99e-22
PB0172.1_Sox1_2
p = 1.07e-21
PH0022.1_Dlx3
p = 6.85e-21
ZKSCAN1
p = 1.45e-28
Gmeb1
p = 3.76e-25
PB0180.1_Sp4_2
p = 1.36e-23
PB0137.1_Irf3_2
p = 1.81e-21
Zfp335
p = 2.84e-21
de novo Motifs
Supplementary Figure 9: Transcription factor binding motif enrichment analysis for hypomethylated promoters in window-based long-read clocks. Motifs were identified using the Hypergeometric Optimization of Motif EnRichment (HOMER) software tool. For each clock, top five known and top five de novo motifs are shown, except for instances when fewer than five significant motifs were identified.

### Slide 10
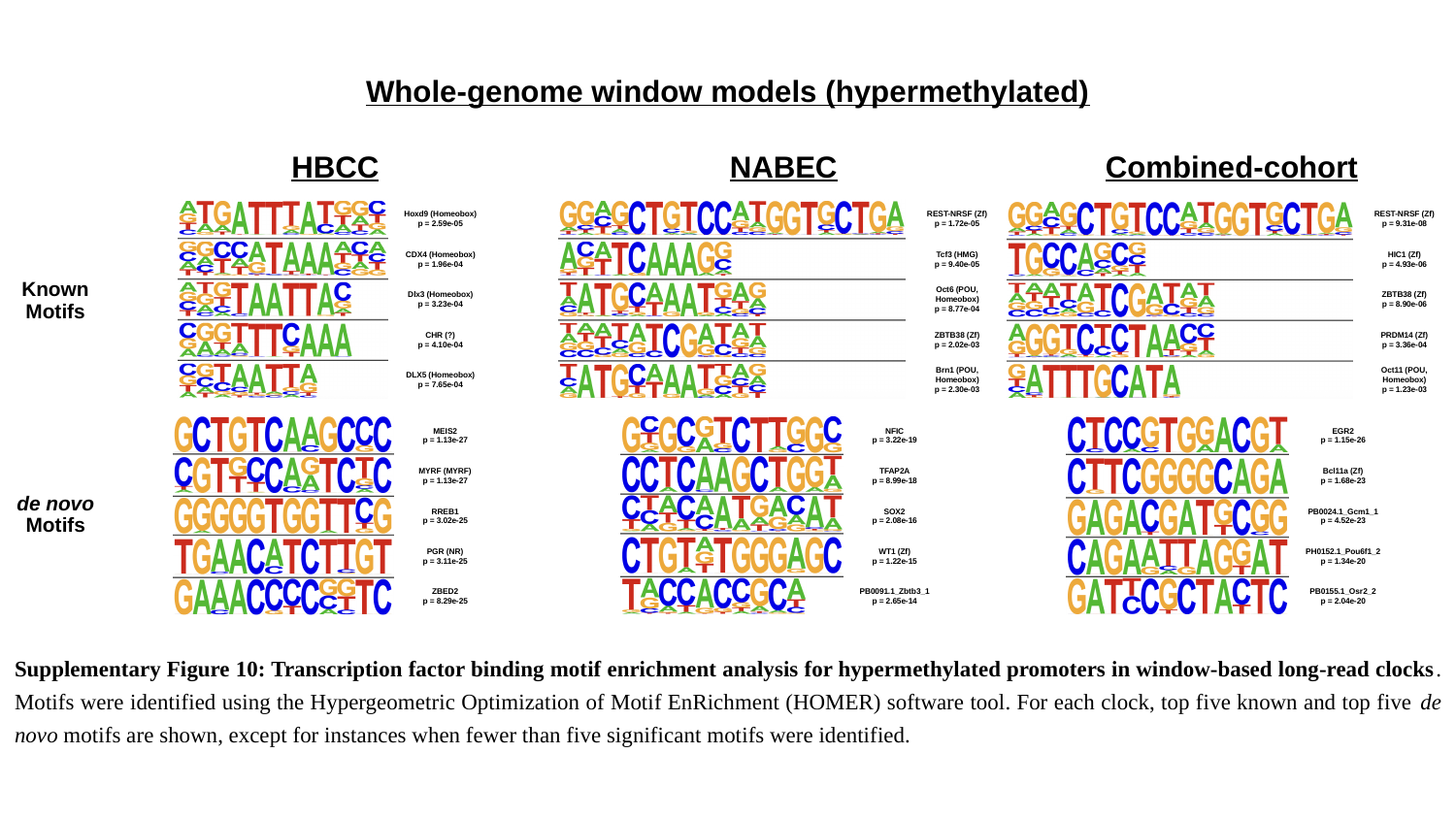

Whole-genome window models (hypermethylated)
HBCC
NABEC
Combined-cohort
Hoxd9 (Homeobox)
p = 2.59e-05
CDX4 (Homeobox)
p = 1.96e-04
Dlx3 (Homeobox)
p = 3.23e-04
CHR (?)
p = 4.10e-04
DLX5 (Homeobox)
p = 7.65e-04
REST-NRSF (Zf)
p = 1.72e-05
Tcf3 (HMG)
p = 9.40e-05
Oct6 (POU, Homeobox)
p = 8.77e-04
ZBTB38 (Zf)
p = 2.02e-03
Brn1 (POU, Homeobox)
p = 2.30e-03
REST-NRSF (Zf)
p = 9.31e-08
HIC1 (Zf)
p = 4.93e-06
ZBTB38 (Zf)
p = 8.90e-06
PRDM14 (Zf)
p = 3.36e-04
Oct11 (POU, Homeobox)
p = 1.23e-03
Known Motifs
MEIS2
p = 1.13e-27
MYRF (MYRF)
p = 1.13e-27
RREB1
p = 3.02e-25
PGR (NR)
p = 3.11e-25
ZBED2
p = 8.29e-25
NFIC
p = 3.22e-19
TFAP2A
p = 8.99e-18
SOX2
p = 2.08e-16
WT1 (Zf)
p = 1.22e-15
PB0091.1_Zbtb3_1
p = 2.65e-14
EGR2
p = 1.15e-26
Bcl11a (Zf)
p = 1.68e-23
PB0024.1_Gcm1_1
p = 4.52e-23
PH0152.1_Pou6f1_2
p = 1.34e-20
PB0155.1_Osr2_2
p = 2.04e-20
de novo Motifs
Supplementary Figure 10: Transcription factor binding motif enrichment analysis for hypermethylated promoters in window-based long-read clocks. Motifs were identified using the Hypergeometric Optimization of Motif EnRichment (HOMER) software tool. For each clock, top five known and top five de novo motifs are shown, except for instances when fewer than five significant motifs were identified.

### Slide 11
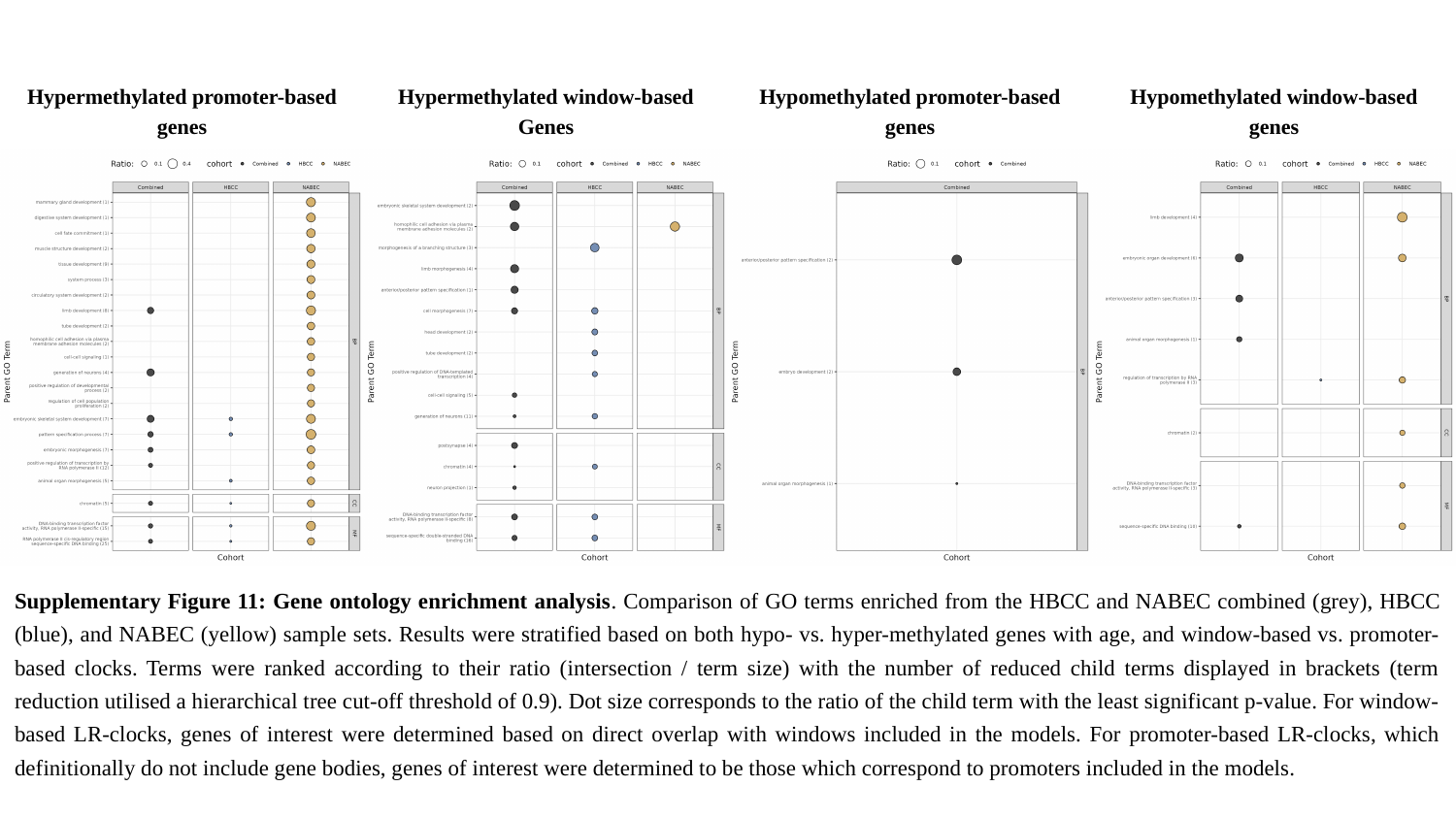

Hypermethylated promoter-based genes
Hypermethylated window-based Genes
Hypomethylated promoter-based genes
Hypomethylated window-based genes
Supplementary Figure 11: Gene ontology enrichment analysis. Comparison of GO terms enriched from the HBCC and NABEC combined (grey), HBCC (blue), and NABEC (yellow) sample sets. Results were stratified based on both hypo- vs. hyper-methylated genes with age, and window-based vs. promoter-based clocks. Terms were ranked according to their ratio (intersection / term size) with the number of reduced child terms displayed in brackets (term reduction utilised a hierarchical tree cut-off threshold of 0.9). Dot size corresponds to the ratio of the child term with the least significant p-value. For window-based LR-clocks, genes of interest were determined based on direct overlap with windows included in the models. For promoter-based LR-clocks, which definitionally do not include gene bodies, genes of interest were determined to be those which correspond to promoters included in the models.

### Slide 12
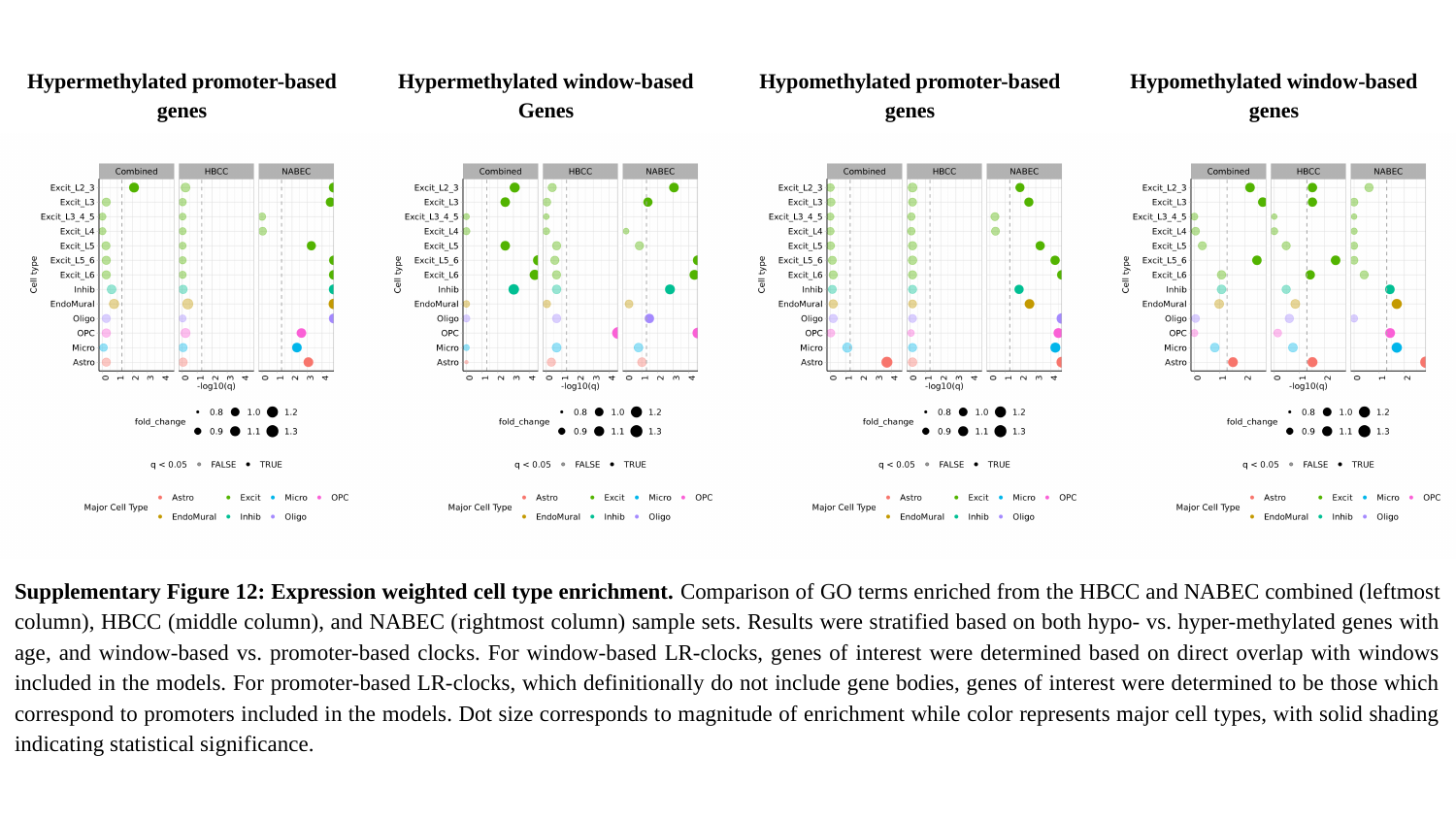

Hypermethylated promoter-based genes
Hypermethylated window-based Genes
Hypomethylated promoter-based genes
Hypomethylated window-based genes
Supplementary Figure 12: Expression weighted cell type enrichment. Comparison of GO terms enriched from the HBCC and NABEC combined (leftmost column), HBCC (middle column), and NABEC (rightmost column) sample sets. Results were stratified based on both hypo- vs. hyper-methylated genes with age, and window-based vs. promoter-based clocks. For window-based LR-clocks, genes of interest were determined based on direct overlap with windows included in the models. For promoter-based LR-clocks, which definitionally do not include gene bodies, genes of interest were determined to be those which correspond to promoters included in the models. Dot size corresponds to magnitude of enrichment while color represents major cell types, with solid shading indicating statistical significance.

### Slide 13
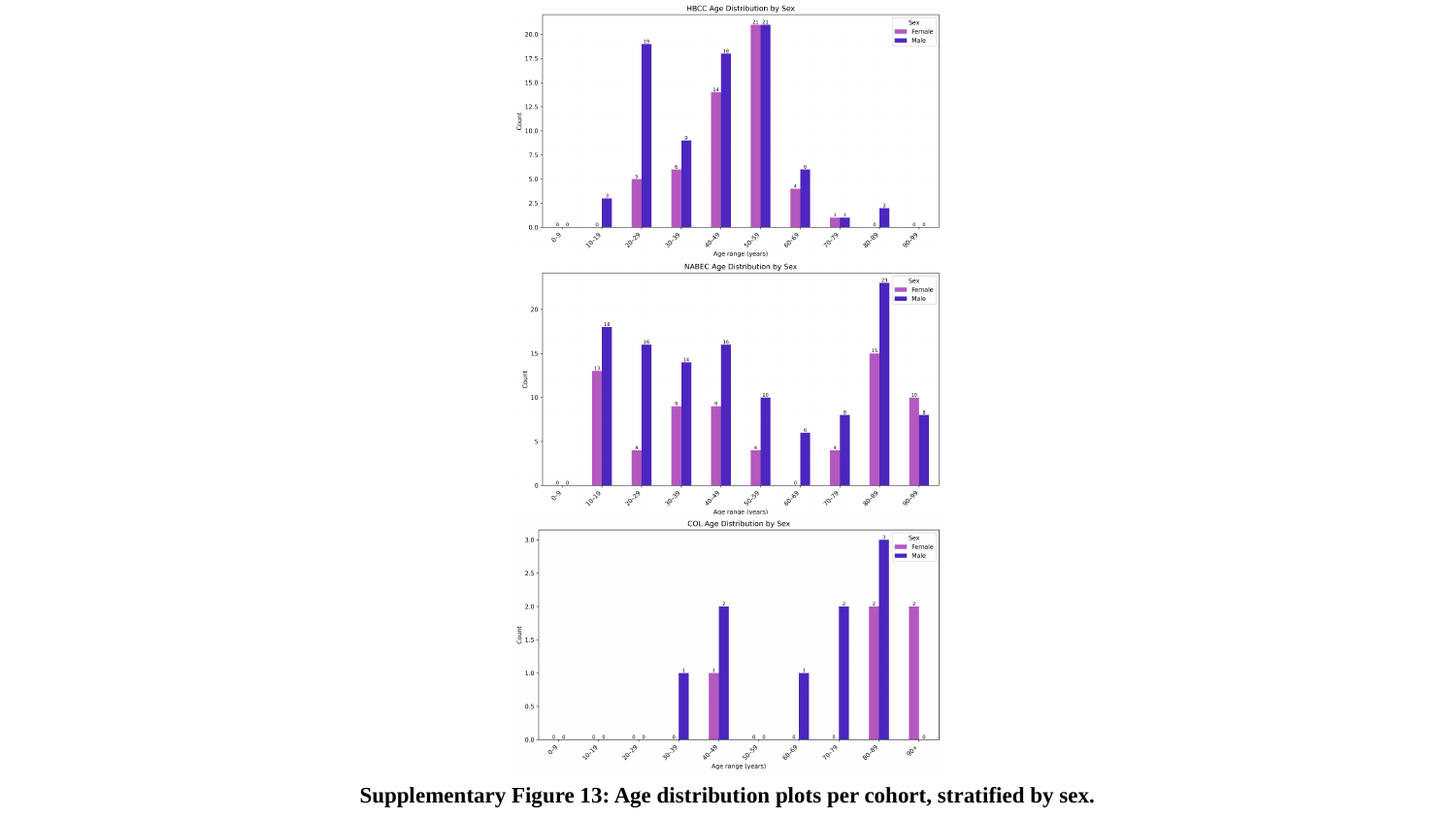

Supplementary Figure 13: Age distribution plots per cohort, stratified by sex.
